# A novel porcine model of CLN3 Batten disease recapitulates clinical phenotypes

**DOI:** 10.1101/2022.10.07.511360

**Authors:** Vicki J. Swier, Katherine A. White, Tyler B. Johnson, Xiaojun Wang, Jimin Han, David A. Pearce, Ruchira Singh, Christopher S. Rogers, Jon J. Brudvig, Jill M. Weimer

## Abstract

Mouse models of CLN3 Batten disease, a rare lysosomal storage disorder with no cure, have improved our understanding of CLN3 biology and therapeutics through their ease of use and a consistent display of cellular pathology. However, the translatability of murine models is limited by disparities in anatomy, body size, life span, and inconsistent, subtle behavior deficits that can be difficult to detect in CLN3 mutant mouse models, limiting their utility in preclinical studies. Here we present a longitudinal characterization of a novel miniswine model of CLN3 disease that recapitulates the most common human pathogenic variant, an exon 7-8 deletion (*CLN3*^*Δex7/8*^). Progressive pathology and neuron loss is observed in various regions of the *CLN3*^*Δex7/8*^ miniswine brain and retina. Additionally, mutant miniswine present with vision impairment and motor abnormalities, similar to deficits seen in human patients. Taken together, the *CLN3*^*Δex7/8*^ miniswine model shows consistent and progressive Batten disease pathology and behavioral impairment mirroring clinical presentation, demonstrating its value in studying the role of CLN3 and safety/efficacy of novel disease modifying therapeutics.

## Introduction

Batten disease [neuronal ceroid lipofuscinoses (NCL)] is a family of 14 different autosomal recessive, pediatric, neurodegenerative disorders that severely reduces quality of life and leads to premature death^1^. The CLN3 subtype (resulting from mutations in the *CLN3* gene), the most common form of Batten disease in the United States and Europe^2^, has a juvenile onset of symptoms between 4 and 7 years of life^3^. Symptoms typically initiate with visual impairment (with loss of photoreceptors), followed by cognitive decline, loss of motor function (including impaired balance and shuffling gate), seizures, and premature death by the second or third decade^4-8^. Pathologically, accumulation of lysosomal storage material, glial activation and neuronal degeneration are key hallmarks of the disease^9^. More than 90 different pathogenic variants have been identified in the *CLN3* gene^10^, the most common being a 966 bp deletion in exons 7 and 8 (Δex7/8)^11^, which occurs in 73% of CLN3 disease patients (85% of CLN3 alleles)^12^.

Multiple mouse models of CLN3 disease have been developed, including *Cln3*^*-/-*13^ knockout models, *Cln3*^*LacZ/LacZ*^ knock-in models^14^, and *Cln3*^*Δex7/8*^ knock-in models with the common human mutation. These *Cln3*^*Δex7/8*^ mice develop classic autofluorescent storage material accumulations, mitochondrial ATP synthase subunit c-reactive deposits (SubC), and microglial and astrocytic activation throughout the brain^9,15,16^. *Cln3*^*Δex7/8*^ mice also demonstrate motor declines evident in gait and coordination tests^9,17-20^ and loss of b-wave retinal function as assessed by electroretinography^21-24^, which has also been documented in *Cln3*^*-/-*^ mice^25^. However, these behavioral phenotypes are subtle and often difficult to recapitulate across laboratories^17,19,20,26^, and *Cln3*^*Δex7/8*^ mice have a variable lifespan^9^, indicating that CLN3 mutation in mouse models lack crucial elements of translatability. Moreover, slight variations in breeding strategies (e.g., background strain), behavior testing paradigms, and/or animal husbandry and environmental enrichment can impact consistency in reported outcomes, including biofluid based biomarkers, in these CLN3 mutant mouse lines^27,28^.

To overcome some of the limitations in the murine models of CLN3 disease, we engineered a novel *CLN3*^*Δex7/8*^ miniswine model. Transgenic swine provide a powerful, translational tool for modeling human diseases that are poorly recapitulated in smaller animal models^29-35^, with their use as a model organism gaining popularity over the last 30 years^36^. More recently, the FDA approved a domesticated pig line as the first ever use of a genetically engineered animal for both food and biomedical/therapeutic purposes^37^. Additionally, swine models are especially useful in neurodegenerative disease modeling due to their gyrencephalic brains, and useful in therapeutic screening, being more similar in size to humans and thus a more human-like physiology and pharmacokinetics. In general, large animal models, such as the CLN5^38^ and CLN6^39,40^ sheep model as well as the CLN2 dog model^41-44^ show more clinical symptoms of Batten disease, and have a longer lifespan allowing for long-term evaluations of disease course and the effectiveness and safety of therapeutics^45^. Given these advantages, our team has developed a novel porcine model of CLN3 disease that replicates histopathological, behavioral, and visual abnormalities experienced by patients with CLN3-Batten disease.

## Results

### Generation of CLN3^Δex7/8^ miniswine, study design, and general observations

rAAV-mediated gene targeting was used to introduce the known disease causing *CLN3* mutation in Yucatan miniswine fetal fibroblasts as previously described^29,32,33,46^. Briefly, male Yucatan fetal fibroblasts were transduced with rAAV carrying a targeting construct designed to replace the endogenous *CLN3* exons 7 and 8 with a neomycin resistance cassette (NeoR) (Supplemental **Figure 1A**), followed by removal of the cassette via cre recombinase–mediated excision. The resulting *CLN3*^*+/Δex7/8*^ fibroblasts were used as nuclear donors for somatic cell nuclear transfer (SCNT). Following SCNT, reconstructed embryos were transferred to recipient miniswine. CLN3-targeted miniswine were born following a 114-day gestation, and heterozygote progenitor miniswine were then bred to expand the colony and generate homozygotes. Successful *CLN3*^*Δex7/8*^ genetic modification was confirmed with PCR and Southern blot (Supplemental **Figure 1B-C**).

A study design schematic is presented in **Figure 1A**, with 29 animals monitored through 36 months of age and 5 animals monitored through 48 months of age. Body score was maintained for all animals over the longitudinal study and no seizures were observed. Social and feeding behavior was considered normal for all animals and no obvious walking impairments were observed by animal staff. Three *CLN3*^*Δex7/8*^ male miniswine developed overt visual dysfunction at 27.8, 38.1, and 42.4 months of age, in which they were observed walking into walls and colliding with unknown structures (outside of home pen) (**Figure 1B**). No survival differences were noted in the 36-to-48-month study period (**Figure 1C**), as Yucatan miniswine can live up to 13-15 years^47^.

**Figure 1:**
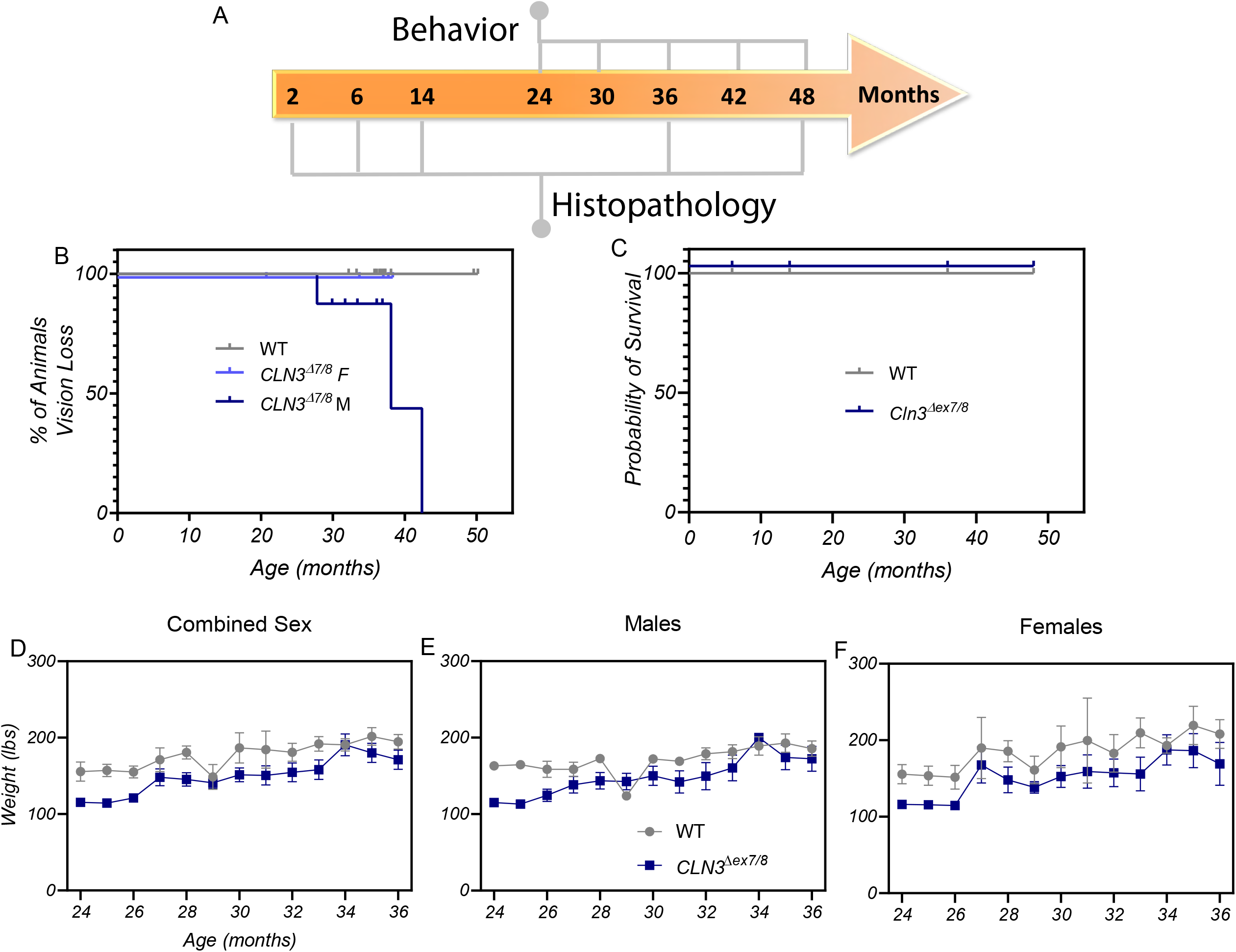
Study design, phenotype monitoring, and weight of miniswine. Study design showing time points for behavior testing and histopathology throughout the study period (A). Percentage of *CLN3*^*Δex7/8*^ and wild-type miniswine with overt vision loss, as documented by animal husbandry staff. Overt vision loss only was observed in only male *CLN3*^*Δex7/8*^ animals (B). Probability of survival of *CLN3*^*Δex7/8*^ and wild-type miniswine, where no early death was detected (C). Monthly weights for combined sexes. *CLN3*^*Δex7/8*^ miniswine weight similar to wild-type (D). Monthly weights for males only. No significant differences are seen between *CLN3*^*Δex7/8*^ males and wild-type males (E). Monthly weights for females. *CLN3*^*Δex7/8*^ females weight similar to wild-type females (F). Mean ± SEM. Mixed-model ANOVA with Sidak’s multiple comparisons.

When examining the weights of all male and female animals at 24 to 36 months of age, *CLN3*^*Δex7/8*^ miniswine weighed similarly to wild-type pigs at all timepoints (**Figure 1D**). When split by sex, male miniswine showed no differences in weight compared to control pigs (**Figure 1E**) and female *CLN3*^*Δex7/8*^ miniswine also weighed similarly to sex matched control pigs (**Figure 1F**).

### CLN3^Δex7/8^ miniswine present with limited cognitive dysfunction but robust motor abnormalities

To determine if *CLN3*^*Δex7/8*^ miniswine recapitulate the cognitive decline displayed in CLN3 disease patients, animals were trained in a simple T-maze as previously described^32,48^. Briefly, the miniswine are allowed to roam the maze for four days for 10 minutes each day in the acclimation phase. Then, the pigs are trained in the acquisition phase to select the food reward arms of the maze for ten-60 second trials on days one and two, during which the reward is placed repeatedly in the same arm of the maze. Lastly, the food is switched to the opposite arm (from the acquisition phase) during the reversal phase to test for the ability to relearn the new task. Wild type and *CLN3*^*Δex7/8*^ miniswine were tested on acquisition and reversal tasks from 24 to 42 months of age, but sustained differences in performance were not detected (**Supplemental Figure 2**). Animals were not tested in the reversal phase at 42 months of age due to poor performance in the acquisition tests at that age (**Supplemental Figure 2C-D**); accuracy on acquisition tests <80%), and tests from 48 months of age are not shown due to low n (n=2/genotype).

To determine if *CLN3*^*Δex7/8*^ miniswine show motor abnormalities, gait was assessed on a pressure-sensor mat from 24 to 48 months of age (to 36 months in females). Significant variables (p<0.05, **Supplemental Table 1**) based on front foot position and pressure were used for synthesis of gait scores based on principal component analysis (PCA), similar to previously described protocols^48,49^. Combined sexes, males, and females were analyzed independently. Based on scores for Principal Component 1 (PC1), *CLN3*^*Δex7/8*^ animals had significantly altered gait compared to wild-type animals at the majority of the time points studied (**Figure 2A-C**). The variables that contributed to the altered gait score in combined sex and male datasets were primarily associated with altered foot strike (footprint size and variability associated with center-of-pressure (COP) trajectory for a single footfall), while female scores were composed largely of spatial variables, such step length variability (**Figure 2D-F**).

**Figure 2:**
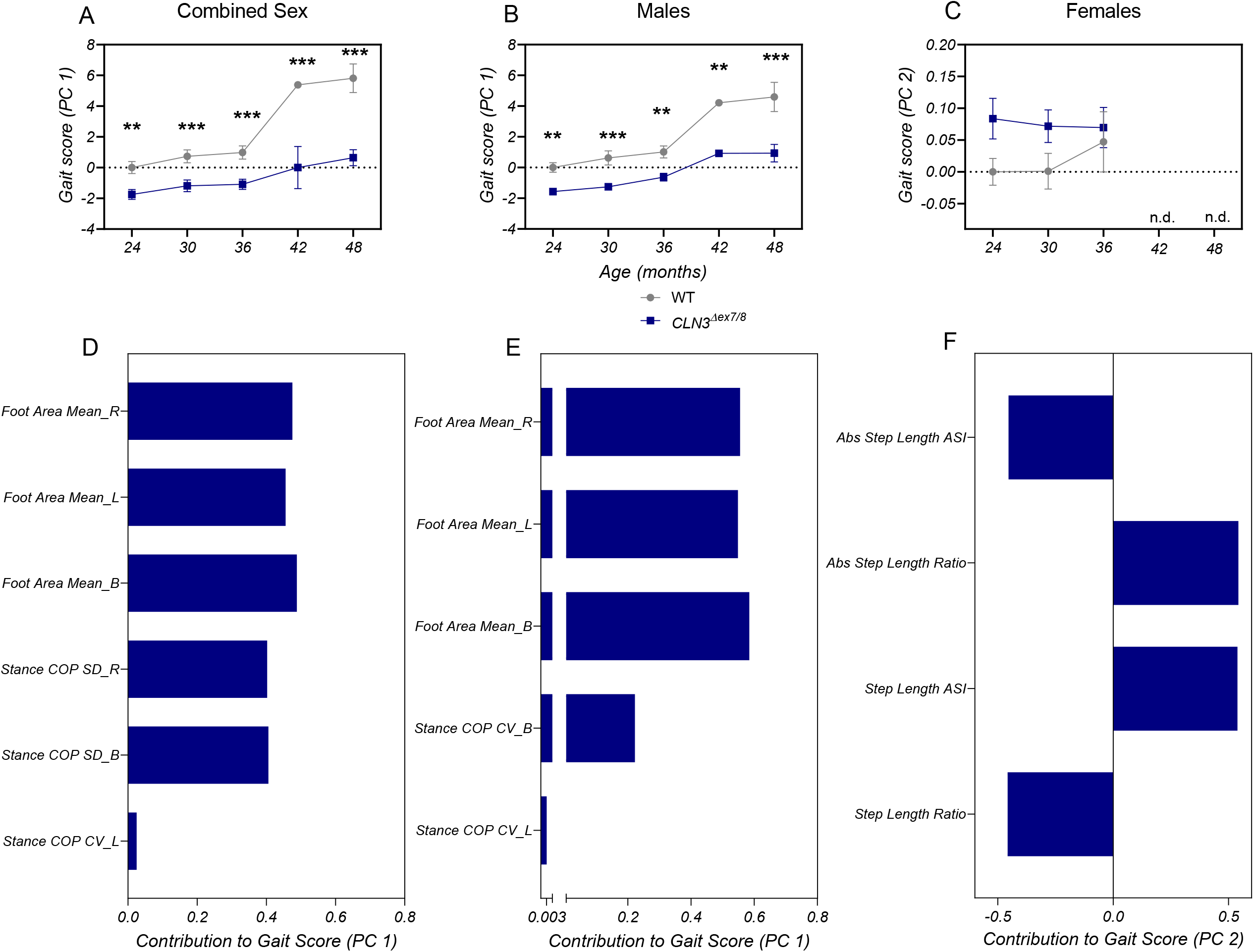
*CLN3*^*Δex7/8*^ animals have a significantly different gait than wild-type. PCA gait score of combined sex (A), male-only (B), and female-only (C) datasets from front feet. *CLN3*^*Δex7/8*^ animals have a significantly different gait than wild-type animals at most time points. Description of contributing variable to gait scores from combined sex (D), male-only (E), and female-only (F) datasets. Two-way ANOVA with uncorrected Fisher’s LSD. **p≤0.01, ***p≤0.001. B=both, R=right, L=left, SD=standard deviation, CV=coefficient of variation.

We next examined how these variables that contributed to the gait scores were indicative of a phenotypic change over time. *CLN3*^*Δex7/8*^ animals, primarily male animals, showed consistently smaller foot area (the area of the foot fall-indicating that the entire foot was not in contact with the ground) than wild-type animals across the time studied (**Figure 3A-B**). Importantly, small foot area was not a result of a smaller body weight (**Figure 1D-F**), indicating differences in *CLN3*^*Δex7/8*^ foot placement upon the mat. Center-of-pressure differences, variables that describe trajectory of a single footfall, were lower in *CLN3*^*Δex7/8*^ combined sex datasets at 30, 36, and 48 months of age, indicating an altered balance phenotype (**Figure 3D)**. Similarly, male *CLN3*^*Δex7/8*^ animals showed lower center-of-pressure variance by 24 months of age (**Figure 3E**). Female *CLN3*^*Δex7/8*^ animals showed less differences over a similar time period, having a more consistent, symmetrical step length, compared to wild-type miniswine (**Figure 3C, F**). Taken together, this data indicates that the front feet of *CLN3*^*Δex7/8*^ animals show altered stepping and center-of-pressure dynamics, similar to the decreased stability observed in individuals with CLN3 disease^50,51^.

**Figure 3:**
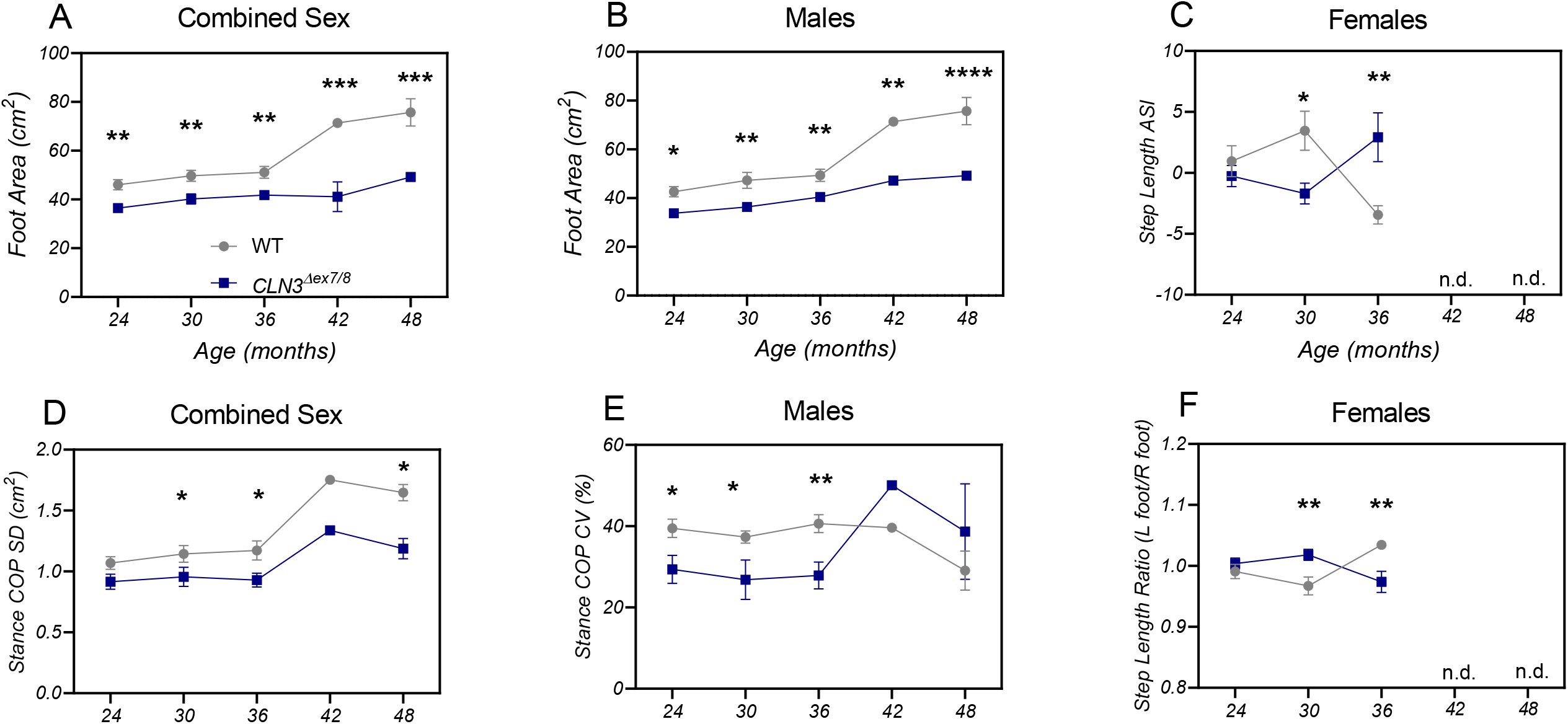
Individual gait variables that contributed to front feet gait score. Foot area for combined sex (A) and male-only (B) datasets. *CLN3*^*Δex7/8*^ miniswine have significantly smaller footfall sizes compare to wild-type at all time points. Stance Center of Pressure (COP) variables for combined sex (D) and male-only (E) datasets. *CLN3*^*Δex7/8*^ miniswine have less variability in stance COP trajectories, indicative of a cautious balancing strategy and more controlled stance. Older *CLN3*^*Δex7/8*^ females show less variability in step length at 30 months of age (C, F). Mean ± SEM. Two-way ANOVA with uncorrected Fisher’s LSD. *p≤0.05, **p≤0.01,***p≤0.001.

The hind feet datasets were also assessed for significant variables for use in the PCAs. Overall, there were few consistent differences between genotypes over the time period studied, though combined male and female *CLN3*^*Δex7/8*^ animals do show altered gait patterns at 24 and 36 months of age, primarily related to greater efficiency of stance in males and less foot pressure in females (**Supplemental Figure 3A-C**). These differences are evident in principal component 1 (PC1) for combined sexes, and principal component 2 (PC2) for males and females. Activity was also assessed via a FitBark activity monitor from 24 to 48 months of age as previously described^31^. No differences in distance traveled, active/rest time, or sleep quality were detected between genotypes (**Supplemental Figure 4A-D**).

### CLN3^Δex7/8^ miniswine exhibit visual decline and photoreceptor loss by 30 months of age

As vision loss is a characteristic phenotype in CLN3 disease, we investigated if *CLN3*^*Δex7/8*^ miniswine showed visual decline. Animals were assessed via flash electroretinography (ERG) from 24 to 48 months of age. By 30 months of age, *CLN3*^*Δex7/8*^ miniswine of both sexes show reduced a-wave (photoreceptor response) and b wave (bipolar cell response) amplitudes in a light adapted, cone-responsive photopic scan, with a- and b-wave responses declining over time (**Figure 4A-F**). Similarly, males also show reduced b-wave amplitudes at 30 months, however reduced a-wave amplitudes are detected later at 42 months of age (**Figure 4B, E)**. No significant differences were detected in females (**Figure 4C, F**). Delayed latencies also arise in *CLN3*^*Δex7/8*^ animals of both sexes at 30 months of age in photopic a- and b-waves (**Figure 4G, J)**. *CLN3*^*Δex7/8*^ males also show delayed latencies at 30 months of age in photopic b-waves, but reduced latencies in a-waves at 36 months of age. Delayed a- and b-wave latencies were detected in *CLN3*^*Δex7/8*^ females at 30 months of age, but this was not sustained (**Figure 4I, L**). When measured in a dark adapted, mixed rod/cone-responsive scotopic ERG, *CLN3*^*Δex7/8*^ miniswine of both sexes show reduced a-wave amplitude at 48 months of age and reduced b-wave amplitude by 42 months age (**Figure 5A, D**). *CLN3*^*Δex7/8*^ males show reduced scotopic a-wave amplitudes at 36 months of age and reduced b-wave amplitudes at 30 months of age (**Figure 5B, E**). No significant differences were found in scotopic a- and b-wave amplitudes in females (**Figure 5C, F**). Significantly delayed latencies in both waveforms arise by 30 months in both sexes of *CLN3*^*Δex7/8*^ animals (**Figure 5G, J**), suggestive of defects in photoreceptor neuronal transmission^52^. Similarly, delayed scotopic a-wave latencies arise at 30 months in *CLN3*^*Δex7/8*^ males and females (**Figure 5H-I**), however slight differences between the sexes are noted in b-wave latencies with delayed latencies at 36 months in males and 30 months in females (**Figure 5K-L**).

**Figure 4:**
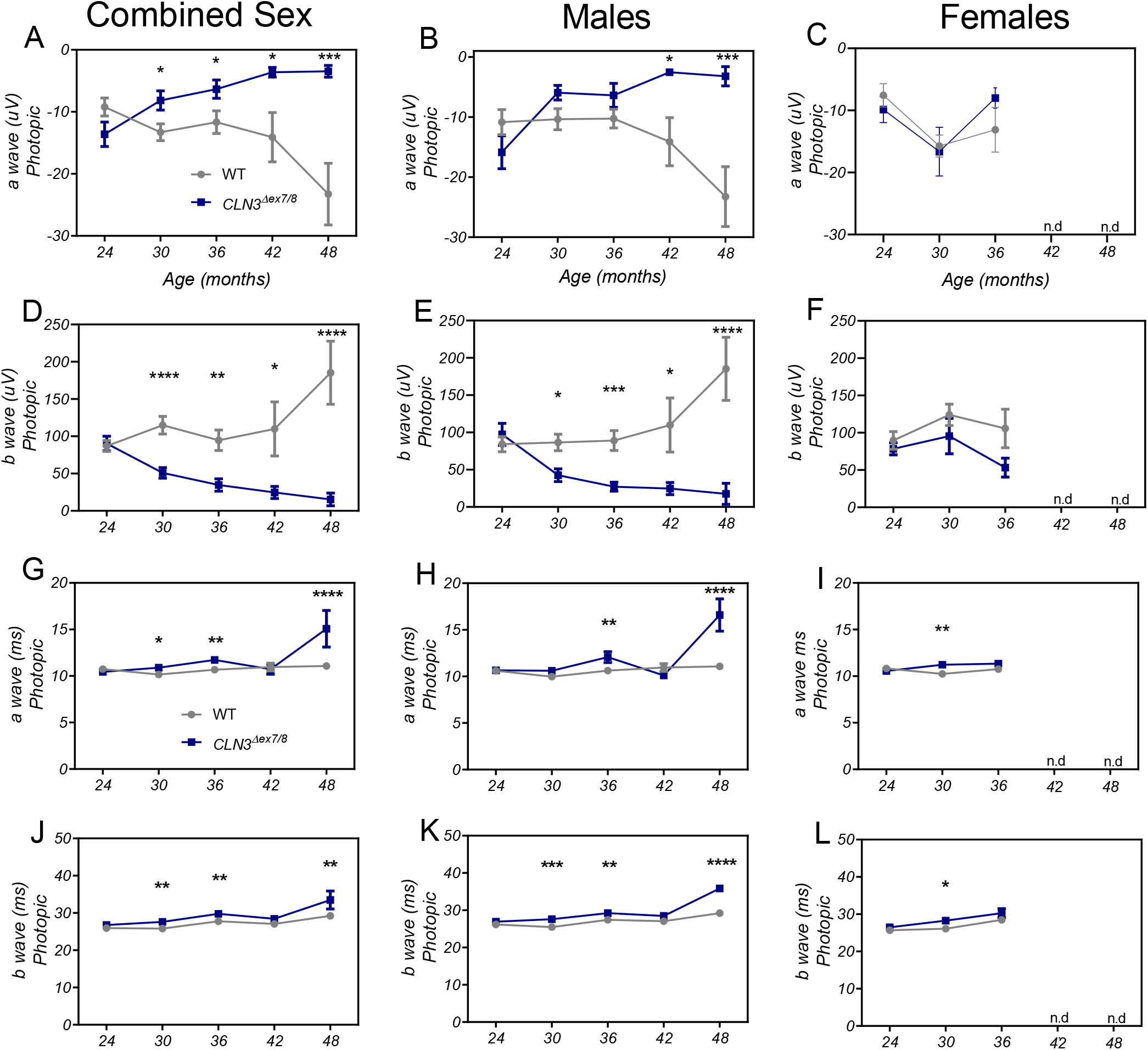
*CLN3*^*Δex7/8*^ animals show photopic visual deficits by 30 months of age as measured by electroretinography (ERG). *CLN3*^*Δe/8*^ miniswine begin to show a reduction in photopic a-wave (A) and b-wave (D) amplitudes at 30 months of age that progressively worsen until the waves are extinguished at 48 months. Photopic a- and b-waves in *CLN3*^*Δex7/8*^ miniswine also show latency (G, J) delays at 30 months. *CLN3*^*Δex7/8*^ males show a-wave amplitude declines at 42 months (B) and b-wave amplitude declines at 30 months (E); Latency delays arise at 36 months in a-wave (H) and slightly earlier at 30 months in b-waves (K). No significant differences in amplitude in female *CLN3*^*Δex7/8*^ miniswine (C, F), and transient latency delays at 30 months in both a- and b-waves (I,L).

**Figure 5:**
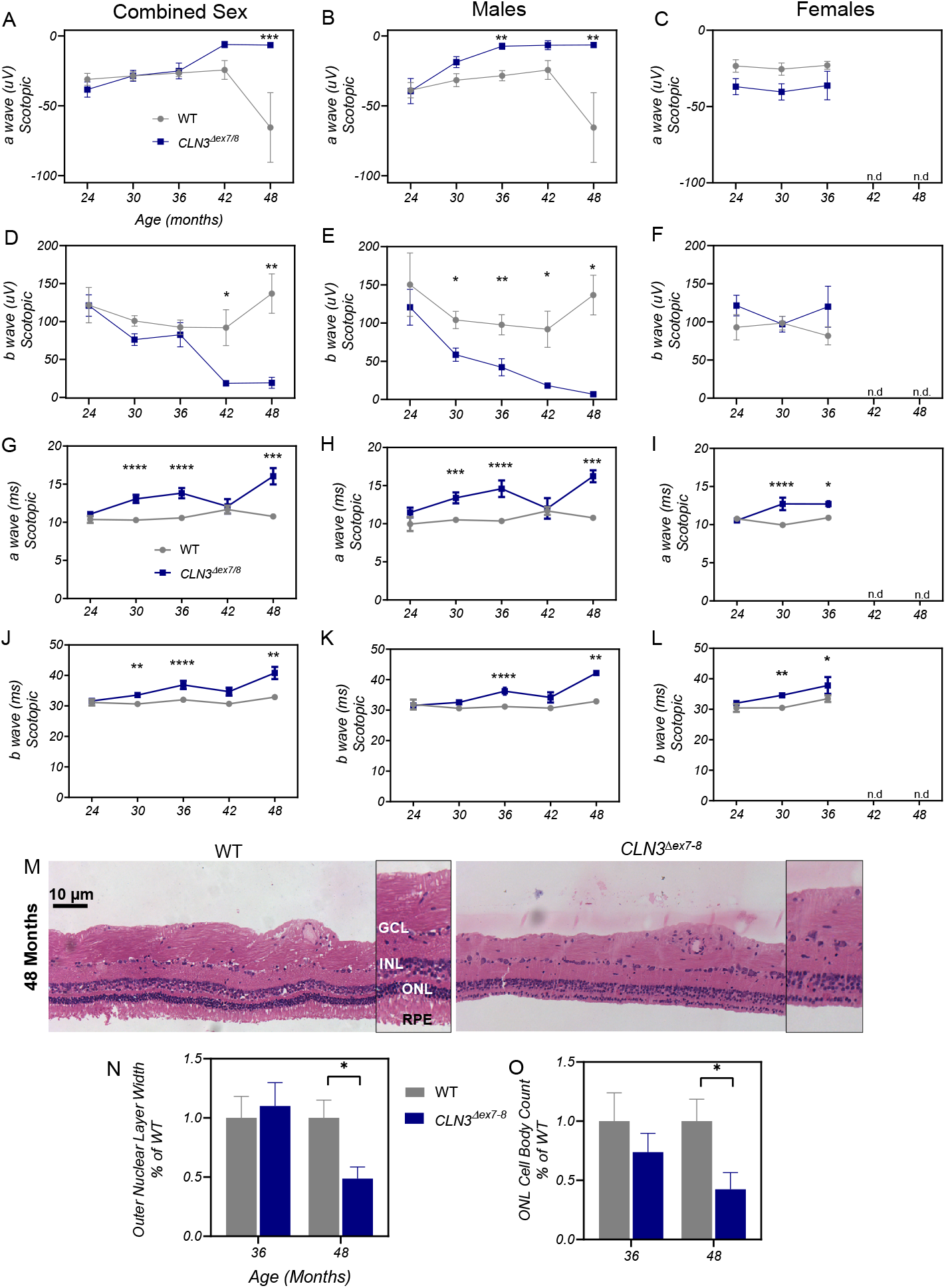
*CLN3*^*Δex7/8*^ animals show scotopic visual deficits by 30 months of age as measured by electroretinography (ERG). Scotopic a-wave (A) and b-wave (D) amplitudes in *CLN3*^*Δex7/8*^ miniswine begin to decline at 42 months of age and are electronegative at 48 months of age. *CLN3*^*Δex7/8*^ male miniswine show declines in scotopic a-wave (B) and b-wave (E) amplitudes by 36 and 30 months of age, respectively. Delayed scotopic a-wave latencies arise at 30 months of age in all *CLN3*^*Δex7/8*^ miniswine (combined sexes, males and females) (G-I). Delayed scotopic b-wave latencies also arise at 30 months in *CLN3*^*Δex7/8*^ miniswine (combined sexes) and females, however slightly later in *CLN3*^*Δex7/8*^ miniswine males at 36 months (J-L). Mean ± SEM. Two-way ANOVA, Fisher’s LSD. *p≤0.05, **p≤0.01,***p≤0.001, ****p≤0.0001. Retinal thinning was measured in 36- and 48-month-old miniswine (M). Outer nuclear layer atrophy was found in animals at 48 months of age (N) and photoreceptor nuclei loss at 48 months (O). Mean ± SEM. Nested t-tests. *p≤0.05.

Retinas were assessed for photoreceptor loss via H+E stain and outer nuclear layer thickness and cell body measurements from 36 to 48 months of age. *CLN3*^*Δex7/8*^ animals showed loss of photoreceptors at 48 months of age, and no loss of cells in the inner nuclear layer (data not shown; **Figure 5M-O**). Importantly, there is a complete loss of the photoreceptor layer in 48-month-old *CLN3*^*Δex7/8*^ animals in the midperiphery of the retina, indicating end-stage retinopathy by this time point, which correlates well to the greatly reduced (and almost extinguished) ERG amplitudes at 48 months.

### CLN3^Δex7/8^ miniswine display classic Batten disease pathology in several regions of the brain

To determine if *CLN3*^*Δex7/8*^ animals exhibit the pathological hallmarks of CLN3 disease, brains were excised, dissected, and examined for classic Batten disease histopathology longitudinally in several brain regions. Specifically, the somatosensory cortex (SSC), motor cortex (MC), VPM-VPL nucleus of the thalamus, and the CA2-CA3 of the hippocampus were assessed (**Figure 6A**). SubC, a common component of Batten disease associated lysosomal storage material, accumulated in *CLN3*^*Δex7/8*^ miniswine by as early as 1 to 4 days of age (**Supplemental Figure 5**) and persisted at 2-, 6-, 14-, 36-, and 48 months of age in the SSC (**Figure 6B-C**), MC (**Figure 6D-E**), CA2-CA3 (**Figure 6F-G**) and VPM-VPL (**Figure 6H-I**).

**Figure 6:**
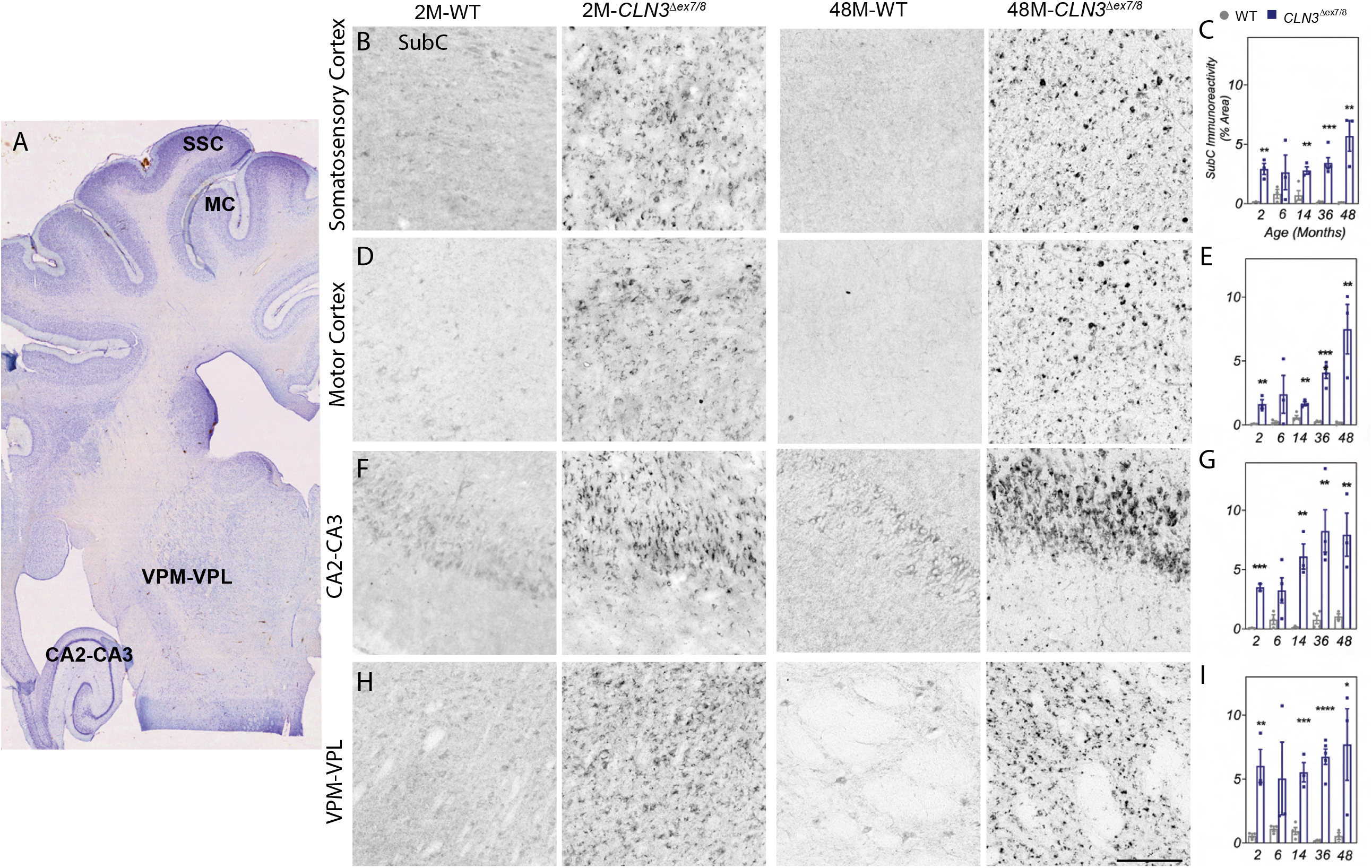
*CLN3*^*Δex7/8*^ animals show mitochondrial ATP synthase subunit c accumulation in several brain regions. Nissl stained coronal section of a miniswine brain indicating the anatomical location of somatosensory cortex, motor cortex, VPM-VPL of thalamus, and CA2-CA3 of hippocampus (A). Subunit C accumulation was evident beginning at 2 months of age in the somatosensory cortex (B-C), motor cortex (D-E), CA2-CA3 (F-G), and VPM/VPL (H-I) of *CLN3*^*Δex7/8*^ miniswine. Accumulation was persistent throughout all time points measured. Mean ± SEM, unpaired t-test, *p≤0.05, **p≤0.01, ***p≤0.001, ****p≤0.0001. Scale bar=200 μm.

In most forms of Batten disease, lysosomal storage and neuronal dysfunction are believed to initiate a neuroinflammatory cascade that results in the activation of glial cells. Iba1+ microglia were examined for their activation status by measuring soma size as previously described^53^. Within the cortex, the soma area was significantly enlarged in the SSC of 36-month-old *CLN3*^*Δex7/8*^ animals (**Figure 7A-B**) and in the MC of 14 and 36-month-old *CLN3*^*Δex7/8*^ animals (**Figure 7C-D**). Surprisingly, there was no microglial reactivity detected in the VPM/VPL (**Figure 7E-F**). Astrocyte reactivity was also examined, and although multiple cortical regions were investigated, strong phenotypic differences in GFAP immunoreactivity were only observed in the thalamus. Reactive astrocytes were first identified in the VPM-VPL as early as 2-month-old *CLN3*^*Δex7/8*^ animals and were sustained longitudinally until 36 months **Figure 7G-H**).

**Figure 7:**
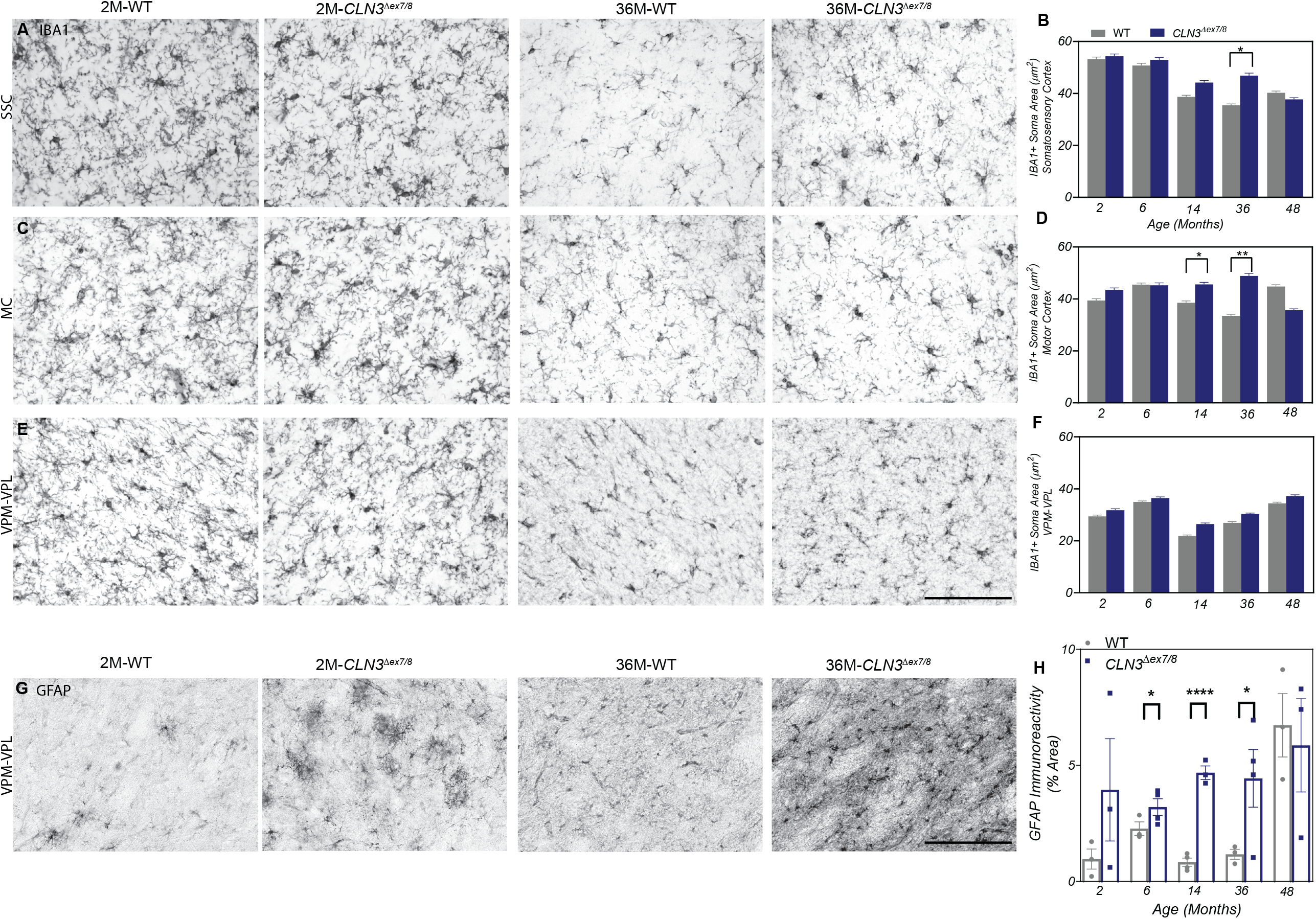
*CLN3*^*Δex7/8*^ miniswine show microglia and astrocyte reactivity in several brain regions. An increase in IBA1+ microglia soma size was evident at 14 to 36 months of age in the somatosensory cortex (A-B) and motor cortex (C-D) of *CLN3*^*Δex7/8*^ animals. Microglial reactivity was not evident in the VPM/VPL (E-F). GFAP+ astrocytosis was evident from 6 to 36 months of age in the VPM/VPL of the thalamus of *CLN3*^*Δex7/8*^ animals (G-H). Mean ± SEM, nested t-test (IBA1) or unpaired t-test (GFAP), *p≤0.05, **p≤0.01, ****p≤0.0001. Scale bars=200 μm.

In Batten disease, neurodegeneration is believed to occur as an end-stage result of neuronal dysfunction and neuroinflammation. To determine whether neurodegeneration was occurring in the brains of *CLN3*^*Δex7/8*^ animals, the thickness of the cortical plate was measured in NeuN labeled sections. No overall cortical atrophy was detected in *CLN3*^*Δex7/8*^ animals from 14 to 48 months (**Supplemental Figure 6**). As particular populations of GABAergic interneurons have been shown to be selectively vulnerable in the cortex of CLN3 patients^54^ and *Cln3*^*-/-*^ mice^13,55^, we also investigated cortical interneuron loss in the miniswine. Calbindin+ neurons were examined in the deepest layers of the cortex (corresponding to layers 5 and/or 6). A significant reduction of calbindin+ interneurons was detected in *CLN3*^*Δex7/8*^ animals at 6 months of age in the SSC and 36 months of age in the MC (**Supplemental Figure 7**), indicating that these cells may be lost progressively in *CLN3*^*Δex7/8*^ animals.

## Discussion

The *CLN3*^*Δex7/8*^ miniswine model recapitulates many human phenotypes of CLN3 disease, including consistent vision loss and gait abnormalities, as well as classic Batten disease pathologies such as storage material accumulation, glial activation, photoreceptor loss and calbindin interneuron loss in the SSC and MC. Importantly, when compared to *CLN3*^*Δex7/8*^ mouse models, the *CLN3*^*Δex7/8*^ miniswine model shows a more severe and consistent phenotype in retinal cell loss, vision loss, and gait abnormalities.

In patients with CLN3 disease, vision loss is one of the first diagnosed symptoms, with the onset of visual failure occurring at a mean age of 6.4 years, with mean age of diagnosis at 8.4 years of age^56,57^. In the *CLN3*^*Δex7/8*^ miniswine, vision loss was first observed in a male miniswine at approximately 28 months of age, which when scaled proportionally to the 15-year lifespan of the Yucatan miniswine model, equates to approximately the age of an older adolescent child. The range of onset does vary in CLN3 patients, males with an earlier onset have been diagnosed with the disease as late as 11.5 years^56^, whereas other studies have found later onsets of 29.7 years (on average) in males and females^58^; which is reflective of the variability in the visual phenotype. In patients, ophthalmic examination may detect retinopathy, maculopathy, a loss of retinal function as indicated by preferential loss of the dark-adapted ERG (scotopic) b-wave amplitude, reduction of scotopic a-waves amplitude, and eventually a reduction of light-adapted ERG (photopic) a- and b-wave amplitudes with a delay in b-wave responses as the disease progresses^57,59-61^. We see similar trends in the *CLN3*^*Δex7/8*^ miniswine, with a significant decrease in scotopic b-wave amplitude present before reductions in scotopic a-waves; and an earlier reduction in photopic a- and b-waves that progressively declines until completely extinguished at later stages of the disease. In contrast, *CLN3*^*Δex7/8*^ mice models show b-wave declines at a much later stage of the disease that is akin to end stage patients^21,23^. Of note, we do see declines in scotopic b-wave amplitudes in the wild type animals as they age, which is reflective of other ERG studies in miniswine, possibly indicating normal aging process of the retina^62^.

Similarly, optical coherence tomography (OCT) imaging in patients with undetectable ERGs shows loss of photoreceptors and marked atrophy of inner and outer nuclear layers of the retina^8,63^. Our *CLN3*^*Δex7/8*^ miniswine model also exhibited a thinning and cell loss in the outer nuclear layer and a complete loss of photoreceptors in the oldest animals monitored, while in contrast, aged *CLN3*^*Δex7/8*^ mouse models have shown only a loss of bipolar cells with no retinal atrophy in the outer nuclear layer^23^. Differences in photoreceptor localization between species may explain the phenotypic differences between *CLN3*^*Δex7/8*^ miniswine and *CLN3*^*Δex7/8*^ mice, as mice lack a macula region of concentrated photoreceptors whereas swine have a macula-like region similar to humans^64^.

We also investigated whether *CLN3*^*Δex7/8*^ miniswine displayed gait abnormalities as a correlate to the loss of motor function, impaired balance, or shuffling gate phenotypes that are common in patients with CLN3^51^. *CLN3*^*Δex7/8*^ male miniswine presented a more cautious gait with smaller footfalls and a controlled stance with lower center of pressure (COP) distance values. COP measurements in gait analysis explain how much an individual moves the directional placement of a footfall’s center of pressure without losing balance, and greater COP distance values indicate a lack of precise, controlled movements^65^. Therefore, lower COP values may be indicative of a careful balancing strategy to compensate for lack of stability. Importantly, reduced variability in COP has been recorded in patients with neurological gait disorders and in patients recovering from strokes as an adaptation to avoid falls and stabilize the gait^66,67^. While *CLN3*^*Δex7/8*^ mouse models present with mild motor coordination phenotypes that are difficult to recapitulate across laboratories, differences in *CLN3*^*Δex7/8*^ miniswine gait were consistent across the time points studied. These differences were primarily male driven, echoing previous reports of sex-specific presentations and disease course in patients and mouse models^20,68^. Interestingly, male *CLN3*^*Δex7/8*^ miniswine had consistently smaller footfalls across the entire time period studied, and as there were no weight differences between genotypes in the male animals, smaller footfall is unlikely to be a result of body size differences. A possible explanation for small footfalls in *CLN3*^*Δex7/8*^ miniswine could be the presence of osteoarthritis, which has been documented in patients with other lysosomal storage disorders such as Mucopolysaccharide Storage Diseases^69,70^, and presents in swine as a convex curvature of the knee that causes the principal digits of the foot to lift off of the ground^71^. This could result in smaller footfalls on a pressure-sensor mat, and more research is needed to understand the nuances of gait abnormalities in swine models of disease.

As observed in patients and *CLN3*^*Δex7/8*^ mouse models, one of the most common markers of Batten disease, ATP synthase subunit C, was found to accumulate throughout multiple brain regions in *CLN3*^*Δex7/8*^ miniswine. Unlike patient and *CLN3*^*Δex7/8*^ mouse models^9,15^, however, astrocytosis was only detected in the VPM/VPL of the thalamus and not in any examined cortical region. One reason could be linked to the increased level of GFAP expression in white matter fibrous astrocytes compared to lower GFAP expression in gray matter protoplasmic astrocytes that has been documented in aging brains^72^, which may indicate the differential upregulation of inflammatory factors in activated white matter astrocytes that are not upregulated in gray matter astrocytes^73^. Other miniswine models of neurodegeneration were also not able to detect GFAP+ astrocytes in cortical gray matter^74^, possibly indicating an even greater differential regulation of GFAP expression in the protoplasmic astrocytes of swine^74^. Additionally, gene expression studies in astrocytes have found species-specific differences between humans and mice, increasing the likelihood that miniswine may also have species specific astrocyte expression profiles that are perhaps not represented through astrocyte labeling that has commonly been used in the Batten disease field^75^.

Although activated astrocytes were not present in the cortex, we did find activated microglia with enlarged somas in the somatosensory and motor cortex in *CLN3*^*Δex7/8*^ miniswine. Activated microglia have been identified in *Cln3* mouse models in multiple cortical regions^76^ and microglia isolated from *CLN3*^*Δex7/8*^ mice have been shown to be primed for activation^77^. Interestingly, two different populations of microglia were isolated from *CLN3*^*Δex7/8*^ mice, one with autofluorescence (AF+) and increased LAMP1 expression, and one population without AF accumulation (AF-) or increased LAMP1; indicating lysosomal disfunction in microglia with accumulation of AF. AF+ microglia also showed increased rates of cell death^78^, indicating that microglia with AF accumulation could be functionally impacted with reduced ability to remove debris, thus contributing to the pathology of *CLN3* disease. As we found SubC accumulation (indicative of AF accumulation) in the cortices of the *CLN3*^*Δex7/8*^ miniswine, this may partially explain the enlarged somas as the microglia are changing to the amoeboid, phagocytizing state to remove excess AF material.

While swine models are touted as more translatable to the human condition, species differences still exist. For example, swine do not present with ulceration of the gastrointestinal tract in stress induced environments, and although the connective tissue of the liver is similar anatomically to humans, swine do not have the same susceptibility to injury-induced cirrhosis of the liver^79^. Additionally, the use of swine and other large animal models of disease are complicated by the lack of species-specific tools, such as antibodies, protocols, behavior equipment, and tissue atlases. Regarding CNS specific disease modeling, brain collection is challenging due to the thicker and denser swine skull, MRI atlases are not readily available for imaging studies, and neurocognitive tests may be difficult to perform or interpret. While there are several caveats and improvements to be made in the use of large animal models of disease, the *CLN3*^*Δex7/8*^ miniswine model is a marked improvement over traditional *CLN3*^*Δex7/8*^ mouse models, with improved size, anatomy, and physiology similarities that are closer to the human condition, along with the presence of robust visual and gait presentations that are typically absent from CLN3 mouse models. *CLN3*^*Δex7/8*^ miniswine therefore hold the promise of improving our understanding of CLN3 disease mechanisms and providing a more relevant setting in which to test therapeutic interventions.

## Acknowledgements

Staff of Precigen Exemplar, including Todd Honkomp, Trisha Smit, Justin Van Kalsbeek, and Cris Van Ginkel for miniswine management and care, Pedro Negrao de assis and Cienna Boss for neurobehavior testing of miniswine, Arlene Drack for advisement on electroretinogram testing and analysis, and Kari Brucker for statistical analysis of IBA1 soma size. Microscopy conducted on microscopes maintained by the Sanford Research Histology and Imaging Core, supported by the Center for Cancer Biology Research CoBRE (NIGMS CoBRE P20GM103548). This project was initiated with an NIH/NINDS SBIR to Exemplar (NS081877).

## Author Contributions

Conceptualization: DAP, CR, JMW; Methodology: VJS, TBJ, XW, JH, RS, CR; Formal Analysis: VJS, KAW, TBJ, XW, JH, JJB; Investigation: VJS, KAW, TBJ, XW, JH, JJB; Writing – Original Draft: VJS, KAW, TBJ, JJB; Writing: Review & Editing: VJS, KAW, TBJ, XW, JH, DAP, RS, CR, JJB, JMW; Visualization: VJS, KAW, TBJ, XW, JH, JJB; Supervision: TBJ, RS, CR, JJB, JMW; Project Administration: VJS, KAW, TBJ, RS, CR, JJB, JMW; Funding Acquisition: DAP, JMW.

## Methods

### Study Approval

All animals were maintained at Exemplar Genetics under an approved IACUC protocol (MRP2015-005).

### Targeting Vector Construction

Genomic DNA was isolated from Yucatan miniswine fetal fibroblasts. A 11.2 kb PCR product that included a region from *CLN3* exon 3 to exon 9 was amplified using a high fidelity polymerase (Platinum Taq High Fidelity, Invitrogen) and *CLN3* primers pCLN3F3 and pCLN3R3 (**Supplemental Table 2**). The PCR product was subcloned into pCR2.1-TOPO (Invitrogen) and sequenced. All DNA sequencing was performed by the University of Iowa DNA Facility. This resulting plasmid served as the template for PCR amplification of the 5′ and 3′ homologous targeting arms, which were subcloned sequentially into a plasmid containing a PGK-NeoR cassette. The primers for the 5′ arm were CLN3 5’armR(EcoRV)2 and CLN3 5’armF(XhoI)2. The primers for the 3′ arm were CLN3 3’armF(HindIII)2 and CLN3 3’armR(HindIII)2. (**Supplemental Table 2**).

### rAAV Production

PCR amplification of a 4.5-kb amplicon from the plasmid described above was achieved by using the following primers: AAVCLN3NeoRF(NotI) and AAVCLN3NeoRR2. This product was subcloned into the rAAV proviral plasmid, pFBAAV2-CMVP.NpA (obtained from University of Iowa Gene Transfer Vector Core) and grown in Sure2 cells (Stratagene). The rAAV was produced by the University of Iowa Viral Vector Core Facility.

### Fetal Fibroblast Infection and Selection

Passage zero male Yucatan fetal fibroblasts (1.0 × 106) were infected with rAAV carrying the *CLN3* targeting construct. After 24 hours, cells were detached with trypsin and plated on 96-well collagen-coated plates. Selection was initiated 48 hours later with G418 (100 μg/ml). Ten to 13 days later, each infected cell plate was split among three 96-well plates (1 plate for freezing, 1 for propagation, and 1 for immediate PCR screening).

### PCR Screen and Cell Handling

Approximately 40% of wells contained live cell colonies following selection. Cells were subjected to 5 μl lysis buffer (50 mM KCl [Sigma], 1.5 mM MgCl2 [Sigma], 10 mM Tris-Cl [RPI Corp], pH 8.5, 0.5% Nonidet P40 [Amresco], 0.5% Tween [RPI Corp], 400 μg/ml Proteinase K [Qiagen]) (75) and incubated at 65°C for 30 minutes, followed by 95°C for 10 minutes. Primers Screen R (NeoR), pCLN32PCRF1 were used to PCR amplify 2 μl of lysate with the following conditions: 2 minutes at 94°C, 30 cycles of 94°C for 20 seconds, 58°C for 20 seconds, and 68°C for 4.5 minutes, and finally 68°C for 3 minutes. The expected product for the targeted allele was a 1.8-kb. The PCR-positive cells were grown to 100% confluence and either infected with rAAV-Cre or expanded for the purpose of DNA isolation.

### Excision of the NeoR Cassette

PCR-positive fetal fibroblast cells were infected with rAAV-CMV-Cre. Three to 6 days later, 90% of the infected cells were frozen, and the remaining cells were propagated for PCR characterization. Cells were lysed in 5 μl lysis buffer as described above, and excision of the selectable marker was detected by PCR using primers pCLN32PCRF1 and pCLN33PCRR18 with the following conditions: 2 minutes at 94°C, 35 cycles of 94°C for 20 seconds, 58°C for 20 seconds, 68°C for 4 minutes, and finally 68°C for 7 minutes.

### Southern Blot

To validate PCR-positive cell lines, genomic DNA was isolated (Gentra, Qiagen) from fibroblasts grown on the propagation culture dishes. Two to 10 ng of genomic DNA was whole genome amplified (Repli-G, Qiagen) and digested with AflII and SspI. Following gel electrophoresis, samples were transferred to a positively charged nylon membrane (Roche Diagnostics) by using an alkaline transfer procedure. The membrane was briefly rinsed in 5× SSC, completely dried, and subjected to UV crosslinking. The DNA probes for *CLN3* and NeoR were produced by PCR amplification using the following primers: pCLN3probeF2/pCLN3probeR3 and NeoR-F/NeoR-R, respectively. Probes were labeled with α-32P by random priming using Prime-a-Gene Labeling System (Promega), and the radioactive probes were purified using CHROMA SPIN+TE-100 columns (Clontech). Membranes were prehybridized in Rapid-hyb Buffer (GE Healthcare) for 30 minutes at 65°C; then, 25 μl of α-32P–labeled probe was added and hybridization proceeded at 65°C for 2 hours. The membrane was washed in 2× SSC and 0.1% SDS once at room temperature for 20 minutes and in 0.1× SSC and 0.1%SDS at 65°C 3 times for 15 minutes each. For confirming animal genotype, high–molecular weight genomic DNA was isolated from miniswine umbilicus. The remaining steps were performed as described above.

### Somatic Cell Nuclear Transfer

Nuclear transfer was performed by Trans Ova Genetics (Sioux Center, Iowa, USA) as previously described^80^. Embryo transfer was performed at Exemplar Genetics. Briefly, reconstructed oocytes were transferred into synchronized post-pubertal domestic gilts on the first day of standing estrus. Recipient gilts were pre-anesthetized with i.v. propofol (0.5–5 mg/kg), and anesthesia was maintained with inhaled isoflurane (3%–5% in oxygen via face mask). Following a midline incision to access the uterus, reconstructed embryos were transferred into the oviduct at the ampullary-isthmus junction. Intra- and postoperative analgesia was provided by i.m. injection of flunixin meglumine (2.2 mg/kg). Recipient animals were checked for pregnancy by abdominal ultrasound after day 21 and throughout gestation.

### Phenotype Monitoring

Wild-type and *CLN3*^*Δex7/8*^ miniswine were monitored from 24 to 48 months of age. Twenty-nine animals were monitored through 36 months of age (wild-type: 9 male/6 female; *CLN3*^*Δex7/8*^: 8 male/6 female) and 5 animals monitored through 48 months of age (wild-type: 2 male; *CLN3*^*Δex7/8*^: 2 male/1 female). Throughout the study, animal husbandry staff monitored body score (score of 1-5 indicating the health by visually inspecting the accumulation of fat around the animal; 1 is emaciated/unhealthy to 5 is overweight)^81^, weight, and the development of Batten disease phenotypes such as signs of vision loss (running into pens, walls, gates; loss of visual tracking), seizures/muscle spasms, walking impairments (off-balance, poor posture, hypermetria), and lethargy, abnormal feeding behavior and assessment of social behavior (type and frequency of vocalizations, interactions with pen mates). Age of onset was recorded when abnormal phenotypes were detected.

### FitBark Activity Monitoring

Miniswine were tracked every six months from 24 to 48 months of age over the course of six days using a FitBark2 device (www.fitbark.com), attached on the neck using a common dog or calf collar as previously described^31,48^. Distance traveled, active time, rest time and sleep quality were analyzed with GraphPad Prism 8.0 using a two-way ANOVA, uncorrected Fisher’s LSD post-hoc.

### Simple T-maze

Memory and learning capabilities were monitored using a simple T-maze as previously described^32,48^, with the exception of using dry animal pellets as a reward. Briefly, during the acquisition phases (two days of testing), animals learned which arm a food reward was located in. During the reversal phases (two days of testing), the reward was moved to the opposite arm and animals had to relearn the location of the reward. Tests were video recorded and tracked with AnyMaze software v4.99 (Stoelting Co. Wood Dale, IL), and a blinded observer watched the T-maze videos and recorded the arm chosen by each animal during each phase. The number of correct responses were analyzed with GraphPad Prism 8.0 using a Mixed-model ANOVA with Sidak’s multiple comparisons.

### Gait assessment

Motor performance was analyzed using a pressure-sensor mat as previously described^48,82^. Briefly, animals were trained to walk on a 4.87×0.6m Zeno Electronic Walkway (ZenoMetrics Peekskill, NY) every six months from 24 months to 48 months of age. Front feet and hind feet data were processed with PKMAS Software ver. 509C1 (Protokinetics LLC, Havertown, PA), and 153 variables were collected per footprint. Five walks per subject were analyzed at each time point, with each walk consisting of eight consecutive footprints. At each time point, t-tests were run on each of the 153 variables within combined sexes, males only and females only to determine which variables were significantly different at p≤0.05. If a variable was significant in at least two timepoints (indicating relevance to a phenotype), that variable was included in a principal component analysis (PCA) implemented in R utilizing the FactorMineR package^83^ to determine which PCAs explained the most variation in the data. Hind gait variables did not reach significance at p≤0.05 within at least two different time points, hence the alpha value was increased to p≤0.1 for the hind foot datasets. T-tests were calculated for each principal component (PC) between wild-type and *CLN3*^*Δex7/8*^ miniswine using pooled time points to determine which PC significantly separated by phenotype at p≤0.05, and the significant PC was used to graph the overall gait score. PC gait scores were standardized to the earliest wild-type average (24 months of age), and thus, the average control score is equal to zero (no deficits). Data were then analyzed in GraphPad Prism 8.0 using a Mixed-model ANOVA with uncorrected Fisher’s LSD.

### Electroretinogram

Miniswine were tested for retinal function using a flash electroretinogram (ERG) every six months from 24 to 48 months of age as previously described^48^. Briefly, animals were anesthetized with 14 mg/kg ketamine (intramuscular) with anesthesia maintained with 1-2% isoflurane. One drop of Tropicamide, Ophthalmic Solution USP, 1% was placed in each eye to cause dilation. Reference electrodes were connected to each ear and a ground electrode to the midline forehead. One drop of Proparacaine Hydrochloride – Ophthalmic Solution, USP, 0.5% was placed in each eye as a local anesthetic and eye speculums were placed inside each eye to fix the eyelids open. The smaller sticky pads of a DTL Plus Electrode (LKC technologies) were attached to the rostral side and larger sticky pads to caudal side of each eye. Each respective DTL electrode (right and left) was connected to the respective extension lead from either the right or left reference electrode, and the ground and both reference electrodes were connected to a RETeval® device (LKC technologies).

The rabbit/minipig, photopic 2 step light adapted protocol was used for each eye and produced an 8.0 cd s/m^2^ flash @ 2.0 Hz flash followed by an 8.0 cd s/m^2^ flicker @ 28.3 Hz. After photopic testing of both eyes, all lights in the room were extinguished, the RETeval® calibrated for dark adaptation, and the animal allowed to dark adapt for 20 minutes. After dark adaptation, the rabbit/minipig, scotopic 4 step protocol was used for each eye. The first step produced a 0.06 cd s/m2 flash @ 0.5 Hz (dark adapted rod only response), followed by an 8.0 cd s/m^2^ flash @ 0.1 Hz (dark adapted mixed rod and cone response), followed by a 25 cd s/m^2^ flash @ 0.05 Hz (dark adapted mixed rod and cone response to higher intensity flash). Raw (unsmoothed) data values were used to calculate amplitudes. a-wave amplitude was recorded as pre-stimulus baseline to a-wave trough, and b-wave amplitude was measured from a-wave trough to the highest waveform peak. Amplitude data from left and right eyes, as well as latency data from the left and right eyes were pooled together for each genotype/time point. a-wave and b-wave amplitudes/peak times for photopic flash responses and scotopic mixed rod/cone responses were analyzed via GraphPad Prism 8.0 using a two-way ANOVA, uncorrected Fisher’s LSD post-hoc.

### Tissue Collection and Processing

Animals were sacrificed with pentobarbital at 2, 6, 14, 36, and 48 months of age for histopathological assessment. One hemisphere of the brain was placed into 10% neutral buffered formalin (∼3 weeks) and subsequently sub-dissected into cortex, hippocampus, and thalamus blocks. Blocks were equilibrated in cryoprotectant solution (30% sucrose in TBSA) at 4°C. Blocks were serial sectioned (50 μm) on a freezing microtome (Leica) and free-floating sections from the somatosensory cortex, motor cortex, VPM-VPL of the thalamus and the hippocampus were placed in 6 well plates for immunohistochemistry.

Eyes were placed in 10% neutral buffered formalin (∼3 weeks) and retinas were removed. Retinal structure was separated from the eyecup and dissected to locate the midperiphery region of the retina. These dissected tissues were further fixed in 4% paraformaldehyde in PBS for 1h at room temperature in glass containers. After secondary fixation, they were washed in PBS and further processed for plastic embedding through steps of dehydration, infiltration, and finally embedding using the Technovit 7100 kit (Electron Microscopy Sciences Cat# 14653). Briefly, retinal tissues were taken through increasing dilutions of ethanol (50% 2h at room temp., 70% overnight at 4C, 80% 1h on ice, 95% 1h on ice, 100% 1h on ice), then 100% acetone 1h on ice for dehydration, and washed a final time in 100% ethanol 1h. Then, for Technovit infiltration retinal tissues were incubated in 1:1 dilution of Technovit Infiltrate/100% ethanol overnight at 4C. The next day, retinal tissue was placed in Technovit Infiltrate for two changes (each 30min) on ice and kept in Technovit Infiltrate overnight at 4C. On the day of embedding, retinal tissues were placed once more in Technovit Infiltrate for 1-2h on ice. Finally, retinal tissues were placed in the polymerization solution for 15min on ice while preparing the plastic molds to embed the tissue. When ready to embed, polymerization solution was pipetted into the embedding cavity of the mold halfway, carefully placing the retinal tissues with forceps in the desired orientation for cutting. Lastly, the rest of the cavity was filled with polymerization solution and a mounting block was positioned, storing at room temp. overnight to allow for polymerization to occur fully. After the plastic was solidified, 3μm-sections were made on the microtome. After sections were dried overnight, they were stained using Multiple Stain Solution (Polysciences, Inc. Cat# 08824-100) for 2min, washed with 70% ethanol, and mounted with permount (Fisher Scientific Cat # SP15-500).

### Immunohistochemistry, Microscopy, and Analysis

Immunostaining for free floating sections followed previous methods ^84^. The following primary antibodies and their dilutions were used: anti-ATP synthase subunit C (Abcam, ab181243; 1:2000), anti-GFAP (Dako, Z0334; 1:16000), anti-IBA1 (BioCare Medical, 290; 1:2000), anti-NeuN (Millipore Sigma, MAB377; 1:2000), and anti-Calbindin (Swant, CB38; 1:2000). Secondary antibodies were biotinylated goat anti-rabbit or biotinylated goat anti-mouse depending on the host of the primary antibody. Secondaries were detected with an avidin-biotin kit (ABC-HRP) (Vector laboratories, CA, USA), followed by incubation in 3,3’-diaminobenzidine (DAB). SubC, GFAP, and NeuN labeled sections were scanned using an Aperio Versa slide scanner (Leica Biosystems, IL, USA) at 20x magnification. At least 3 images were extracted from the somatosensory cortex, motor cortex, hippocampus and VPM-VPL of the thalamus using Aperio ImageScope 12 software (Leica Biosystems, IL, USA), and images were condensed by Adobe Photoshop and split into RGB channels by ImageJ (NIH)^85^. For SubC and GFAP, total percent area was performed with adjusted threshold and analyze particles settings in ImageJ. For NeuN, cortical plate thickness was measured in coronal sections of the somatosensory and motor cortex following methods previously published ^86^. IBA1 and calbindin labeled sections were imaged on a Nikon 90i microscope (Nikon instruments, Inc, NY, USA) at 20x magnification, images condensed with Adobe Photoshop, and split into RGB channels with ImageJ. For IBA1 and calbindin, soma size and number was measured by adjusting the threshold and particle size settings in ImageJ software (NIH).

Retinal sections from each animal were imaged at 10X, 20X, and 40X. The inner and outer nuclear layer were measured in 10X retinal images with ImageJ, measuring the thickness of each layer in triplicates using the line tool in NIS-Elements. The outer nuclear layer thickness was additionally quantified by counting the number of photoreceptor nuclei stacked in the ONL in triplicates from each 20X image (≥5 for each animal). ImageJ software was used to mark nuclei that were counted and images were randomized for both age and sex of the animal prior to quantification to diminish bias. For each image, the count was taken from the middle, left, and right areas of the retina presented in each image, making efforts to stay consistent with each region from image to image.

### Statistics

All data analyses were performed with GraphPad Prism 8.0 or equivalent. Outliers were removed using the ROUT method, Q=1. For each timepoint, unpaired t-tests were used to: compare the mean total percent immunoreactivity of SubC and GFAP; to compare the means for cortical plate thickness; and to compare the mean number of calbindin+ cells between *CLN3*^*Δex7/8*^ and wild-type miniswine. For each timepoint, nested t-tests were used to: compare IBA1+ soma sizes per animal; and retinal layer thickness per animal. Detailed statistical tests are described in the figure legends. *p<0.05, **p<0.01, ***p<0.001, ****p<0.0001. Animal numbers for behavior testing are listed in **Supplemental Table 3**.

