## Supplemental Figures and Tables for "A novel porcine model of CLN3 Batten disease recapitulates clinical phenotypes"

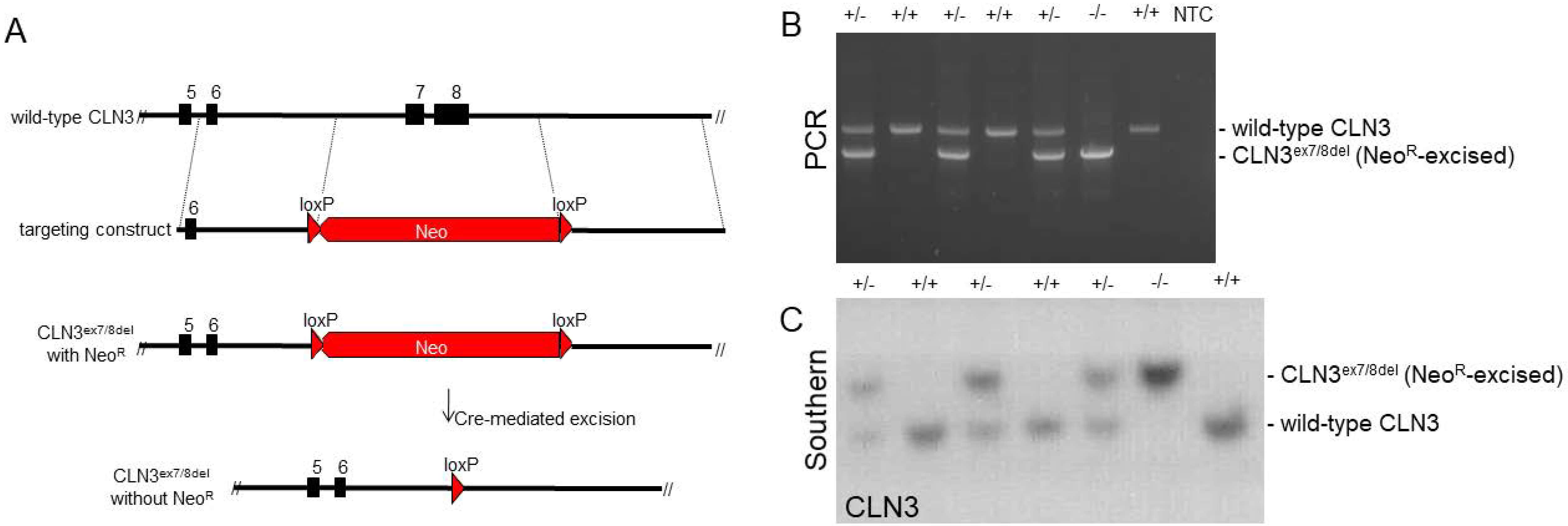

A
5
6
7
8
wild-type CLN3
6
loxP
loxP
targeting construct
Neo
5
6
loxP
loxP
CLN3ex7/8del
with NeoR
Neo
Cre-mediated excision
5
6
loxP
CLN3ex7/8del
without NeoR
B
+/-
+/+
+/-
+/+
+/-
-/-
+/+
NTC
PCR
- wild-type CLN3
- CLN3ex7/8del (NeoR-excised)
C
+/-
+/+
+/-
+/+
+/-
-/-
+/+
Southern
CLN3
- CLN3ex7/8del (NeoR-excised)
- wild-type CLN3

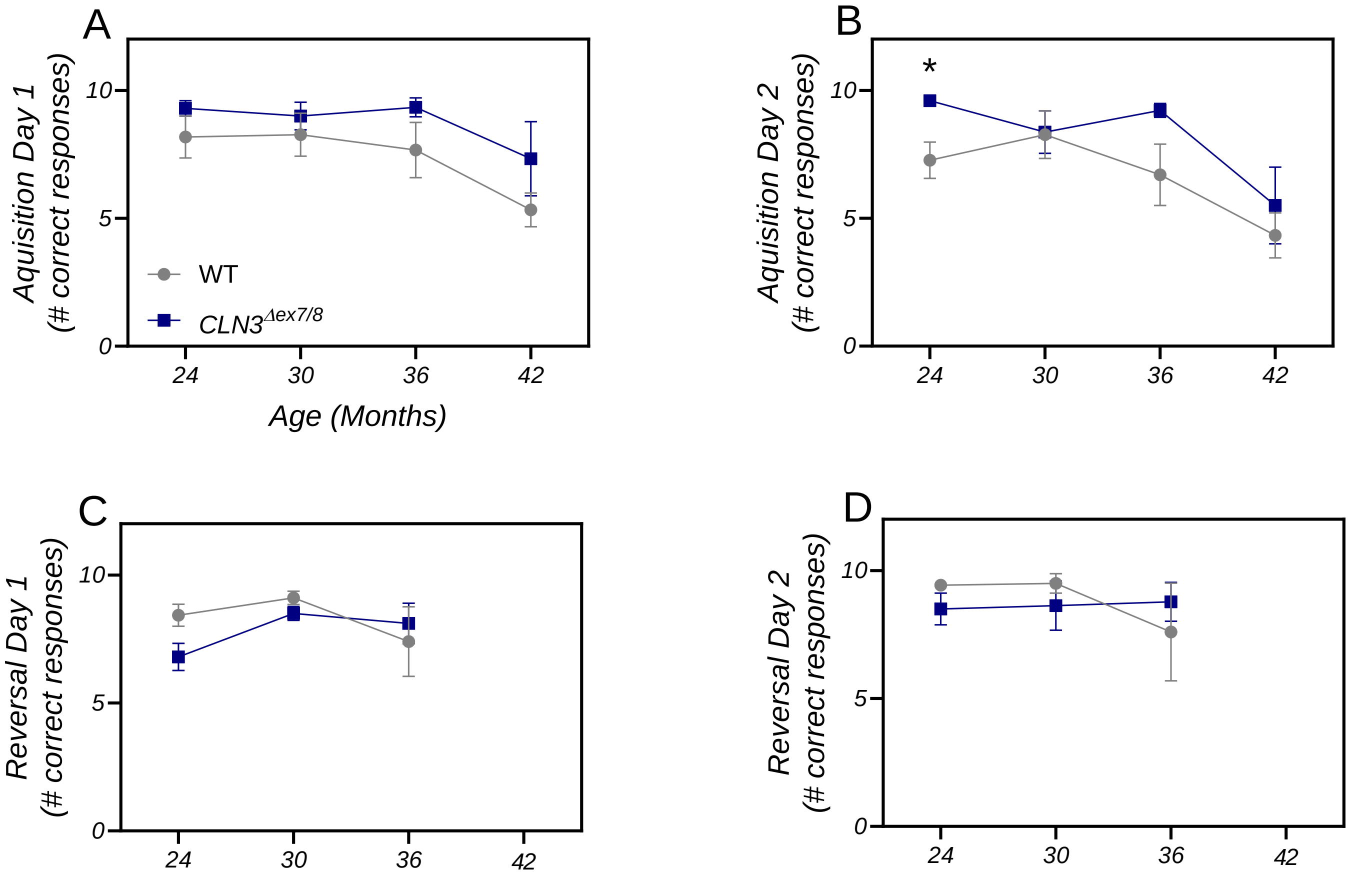

A
Aquisition Day 1
(# correct responses)
10
5
0
WT
CLN3Δex7/8
24
30
36
42
Age (Months)
B
*
Aquisition Day 2
(# correct responses)
10
5
0
24
30
36
42
C
Reversal Day 1
(# correct responses)
10
5
0
24
30
36
42
D
Reversal Day 2
(# correct responses)
10
5
0
24
30
36
42

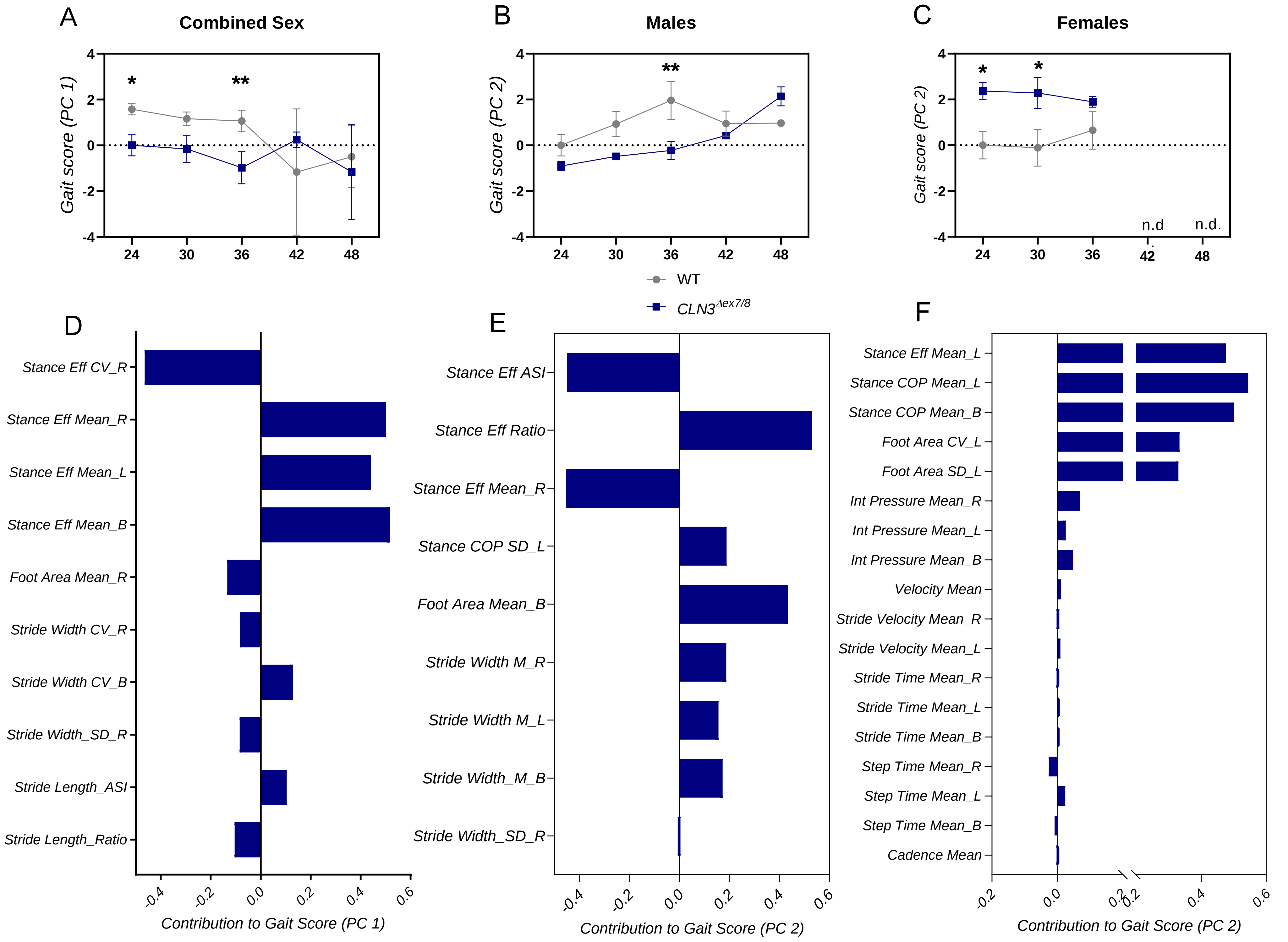

A
Combined Sex
B
Males
C
Females
Gait score (PC 1)
Gait score (PC 2)
*
**
n.d
n.d.
24
30
36
42
48
WT
CLN3Δex7/8
D
E
F
Stance Eff CV_R
Stance Eff Mean_R
Stance Eff Mean_L
Stance Eff Mean_B
Foot Area Mean_R
Stride Width CV_R
Stride Width CV_B
Stride Width_SD_R
Stride Length_ASI
Stride Length_Ratio
Contribution to Gait Score (PC 1)
Stance Eff ASI
Stance Eff Ratio
Stance Eff Mean_R
Stance COP SD_L
Foot Area Mean_B
Stride Width M_R
Stride Width M_L
Stride Width_M_B
Stride Width_SD_R
Contribution to Gait Score (PC 2)
Stance Eff Mean_L
Stance COP Mean_L
Stance COP Mean_B
Foot Area CV_L
Foot Area SD_L
Int Pressure Mean_R
Int Pressure Mean_L
Int Pressure Mean_B
Velocity Mean
Stride Velocity Mean_R
Stride Velocity Mean_L
Stride Time Mean_R
Stride Time Mean_L
Stride Time Mean_B
Step Time Mean_R
Step Time Mean_L
Step Time Mean_B
Cadence Mean
Contribution to Gait Score (PC 2)

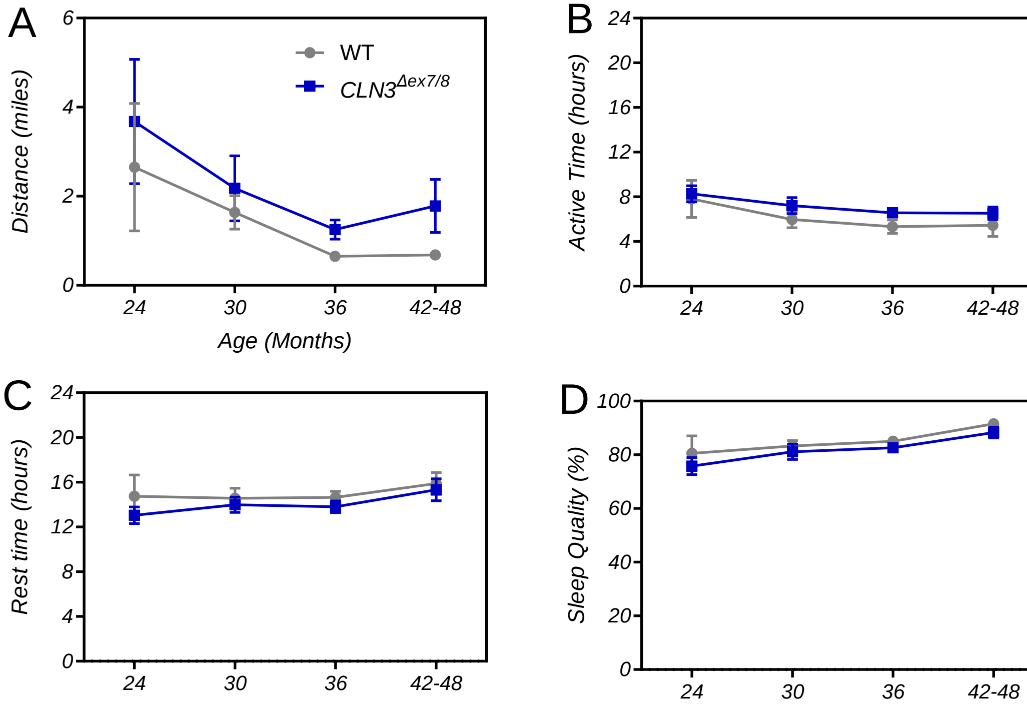

A
B
C
D
WT
$CLN3^{\Delta ex7/8}$
Distance (miles)
Active Time (hours)
Rest time (hours)
Sleep Quality (%)
Age (Months)
24
30
36
42-48
0
2
4
6
8
12
16
20
40
60
80
100

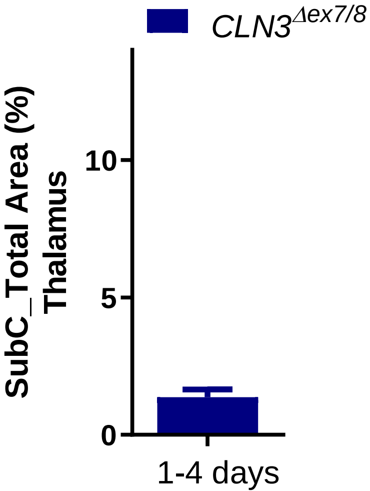

$CLN3^{\Delta ex7/8}$
SubC_Total Area (%)
Thalamus
10
5
0
1-4 days

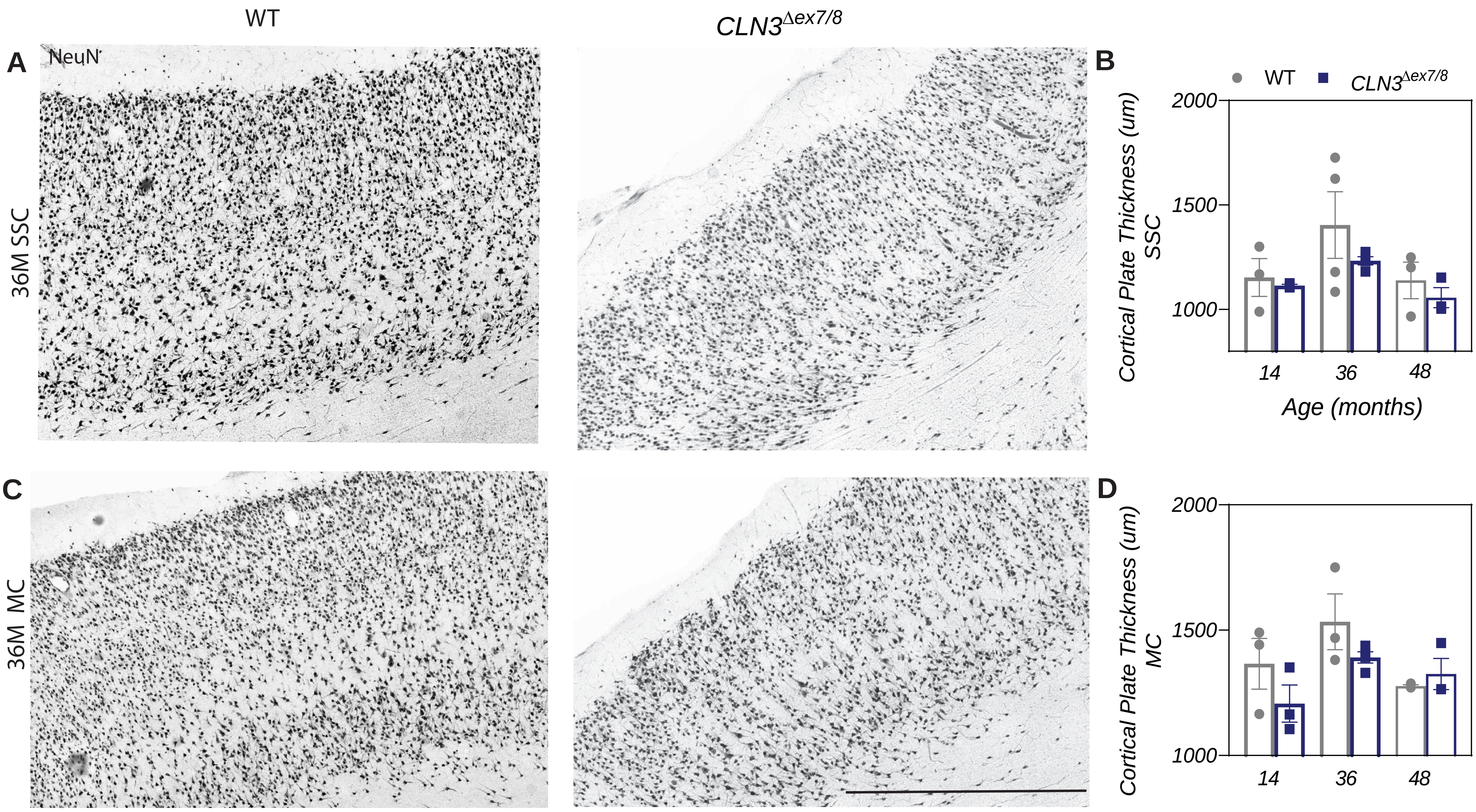

WT
$CLN3^{\Delta ex7/8}$
A
NeuN
36M SSC
B
WT
$CLN3^{\Delta ex7/8}$
Cortical Plate Thickness (um) SSC
2000
1500
1000
14
36
48
Age (months)
C
36M MC
D
Cortical Plate Thickness (um) MC
2000
1500
1000
14
36
48

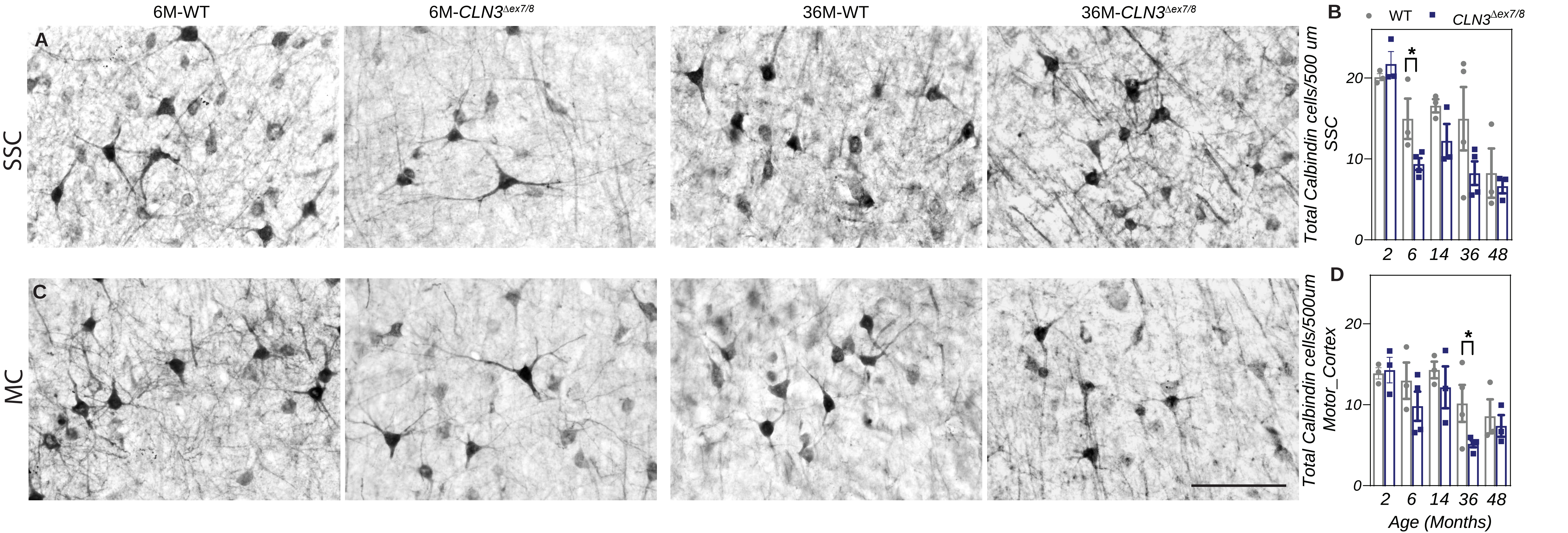

6M-WT
6M-CLN3Δex7/8
36M-WT
36M-CLN3Δex7/8
A
B
C
D
SSC
MC
WT
CLN3Δex7/8
Total Calbindin cells/500 um SSC
Total Calbindin cells/500um Motor_Cortex
Age (Months)
2
6
14
36
48
0
10
20
*

**Supplemental Figure 1: *CLN3$^{\Delta ex7/8}$* founder generation**. Schematic of gene targeting vector used to delete exons 7/8 from porcine CLN3. Porcine CLN3 exons are depicted as black boxes. Neomycin resistance cassette (red) is driven by the phosphoglycerate kinase (PGK) promoter and flanked by loxP sites. Each homology arm is ~1.4 kb in length. Following Cre-mediated excision, a single loxP site remains (A). PCR confirmation of *CLN3$^{\Delta ex7/8}$* recombination in miniswine; WT: lanes 2, 4, 7; Heterozygote: 1, 3, 5; Homozygote: lane 6. Lane 8 is a no template control (B). Southern blot confirmation of CLN3$^{\Delta ex7/8}$ allele in miniswine; WT: lanes 2, 4, 7; Heterozygote: 1, 3, 5; Homozygote: lane 6 (C). When probed for Neo$^{R}$, no signal was detected, as would be expected following Neo$^{R}$ excision (data not shown).

**Supplemental Figure 2: *CLN3$^{\Delta ex7/8}$* miniswine do not show learning deficits.** All animals perform similarly during Acquisition Day 1 (memory test) (A)**,** Acquisition Day 2 (memory test) (B), and Reversal Day 1 (learning test) (C), with a similar number of correct choices. However, at 42 months of age, no animals were able to complete the learning test due to poor performance in the memory tests (<80% accuracy; n.d. represents no data). All animals perform similarly during Reversal Day 2 (learning test), with a similar number of correct choices (D)**.** Mean ± SEM. Mixed-model ANOVA with Sidak's multiple comparisons.

**Supplemental Figure 3**: *CLN3$^{\Delta ex7/8}$* animals have a significantly but subtly different hind gait than wild-type at 24 and 36 months of age. PCA gait score of combined sex (A), male-only (B), and female-only (C) datasets from hind feet. Description of contributing variable to gait scores from combined sex (D), male-only (E), and female-only (F) datasets. Two-way ANOVA with uncorrected Fisher's LSD. *p≤0.05, **p≤0.01.

**Supplemental Figure 4:** *CLN3$^{\Delta ex7/8}$* and wild-type animals do not show any significant differences in home pen activity using a Fitbark activity monitoring device, including total distance travelled (A), time active (B), time resting (C), and sleep quality (D). Mean ± SEM. Two-way ANOVA, Fisher's LSD.

**Supplemental Figure 5**: Subunit C accumulation was evident at 1-4 days in the thalamus of *CLN3*$^{\Delta ex7/8}$ miniswine. Mean ± SEM.

**Supplemental Figure 6:** *CLN3*$^{\Delta ex7/8}$ animals do not show cortical atrophy at 14, 36 or 48 months of age in either the somatosensory cortex (A-B) or motor cortex (C-D). Neurons labeled with NeuN. Mean ± SEM. Unpaired t-tests. Scale bar=700 µm.

**Supplemental Figure 7**: From 6 to 36 months of age, a reduced number of calbindin+ interneurons are found in the somatosensory cortex (A-B) and motor cortex (C-D). Mean ± SEM, unpaired t-test, *p≤0.05, Scale bar=200 µm.

| Combined Sex | | |
|---|---|---|
| Gait variable | p value | Time points |
| Foot Area Mean R | 0.009, 0.02, 0.01, 0.04 | 24M, 30M, 36M, 48M |
| Foot Area Mean L | 0.003, 0.005, 0.02, 0.04 | 24M, 30M, 36M, 48M |
| Foot Area Mean B | 0.002, 0.006, 0.007, 0.04 | 24M, 30M, 36M, 48M |
| Stance COP SD R | 0.01, 0.01 | 30M, 36M |
| Stance COP SD B | 0.02, 0.05 | 36M, 48M |
| Stance COP CV L | 0.002, 0.01 | 36M, 48M |

| Male only | | |
|---|---|---|
| Gait variable | p value | Time points |
| Foot Area Mean R | 0.04, 0.04 | 36M, 48M |
| Foot Area Mean L | 0.01, 0.01, 0.04 | 24M, 30M, 48M |
| Foot AreaMean B | 0.007, 0.01, 0.02, 0.04 | 24M, 30M, 36M, 48M |
| Stance COP CV B | 0.04, 0.05, 0.007 | 24M, 30M, 36M |
| Stance COP CV L | 0.002, 0.01 | 36M, 48M |

| Female only | | |
|---|---|---|
| Gait variable | p value | Time points |
| Abs Step Length ASI | 0.02, 0.01 | 30M, 36M |
| Abs Step Length Ratio | 0.02, 0.01 | 30M, 36M |
| Step Length ASI | 0.02, 0.01 | 30M, 36M |
| Step Length Ratio | 0.02, 0.01 | 30M, 36M |

Supplemental Table 1: Significant gait variables, p value for each unpaired t-test, and time points in which the variables were significantly different between CLN3Δ7/8 and WT animals for combined sexes, males only and females only. SD-standard deviation, CV-coefficient of variation, R-right, L-left, B-both.

| List of CLN3 primers for making targeting construct |
|---|
| pCLN3F3 |
| GTT TAG CTG CTC TTA AAG GTA C |
| pCLN3R3 |
| CTG CTG AGC ATG ACT TAG GA |
| pCLN3seqF9 |
| TGA CTG CAC ACG TGG CAT GCA |
| pCLN3seqR10 |
| GTG GCT CTG GTT CCC AGG TGC |
| pCLN3seqF12 |
| TGG ACC CAG ACC CAA CAC CCA |
| pCLN3seqR15 |
| TAG GGC AGC AGA TGG AGG CCA |
| pCLN3seqF14 |
| TGC TCC TGG CAG ACA TCC TTC |
| pCLN3seqR13 |
| AGC CGT GGA GAC AGA GTT ACA |
| pCLN3seqF11 |
| GCT CCT GGG CCT CTG CAA CAA |
| pCLN3seqF19 |
| GTG AGG AAG TGT CAT GGT CTG |
| pCLN3seqR20 |
| CTA TTG GCA TCC AGC AGG TAG |
| pCLN3Ex6F7 |
| AGC TTC ATC TTG GTC GCC TTC |
| pCLN3Ex7F4 |
| TCT TGG CTA GCA TCT CTT CAG |
| pCLN3Ex8R4 |
| GCT AGC ATC AGG GCA GGG ATA |
| pCLN3Ex9F17 |
| TCC TGT TGC TCA CGT CTC CTG |

| |
|---|
| pCLN3seqR16 |
| CAC AAT GCA GTG AGA CTT CTT |
| pCLN3Ex9R18 |
| CGC TAT TTA TCA GGG GCT GCC |
| pCLN3seqR24 |
| CAG CTA CAG CTC CAA TTG GAC |
| pCLN3seqF21-2 |
| AGG CTC TGA TAG GCC TGT TTG |
| pCLN3seqR25 |
| GTA TAC ATG TAT GTG TAA CTG |
| pCLN3seqF28 |
| TTA GGG CCA GGC CTT GTA GAA |
| pCLN3seqR27 |
| TTG ACT CCT GGC TCA CAC CAA |
| CLN35'armR(EcoRV)2 |
| ATC TGG GAT ATC TGT GGC TGT GGC GTA GGC CTG |
| CLN35'armF(XhoI)2. |
| TGT GAG CTC GAG TGT AGG CCA GTA GCT ACA GCT |
| CLN33'armF(HindIII)2 |
| TGG AGC AAG CTT TTG TTG GCT GTG TTG TAT GGG |
| CLN33'armR(HindIII)2 |
| AAG GGG AAG CTT CGT CTC CTA CCT GGC TTC AAC |
| AAVCLN3NeoRF(NotI) |
| CAC TAG TCG CGG CCG CTA CCA CTG AGC CGC AAT GGG A |
| AAVCLN3NeoRR2 |
| TAG GTC GCA GCG GCC GCT CAG GTC CTG TGT TGT TGT GGC TG |
| AAVF1 |
| CTC TAG CTA TAG TTC TAG TGG |
| AAVR2 |
| GTG GTA TGG CTG ATT ATG ATC |
| Screen R (NeoR), |
| AAG ACA ATA GCA GGC AAC AAC |
| pCLN32PCRF1 |

| |
|---|
| CTT ACC CTT ACT CTG GGT CTG TAG |
| pCLN33PCRR18 |
| TGG TCA GAG AGG TAA AGT AAC |
| pCLN3probeF2 |
| TAA GAA GCC AAT GCT GGA GTT |
| pCLN3probeR3 |
| AAG TAT TCG GCG TGG CCG TGA |
| NeoR-F |
| GCC ATT GAA CAA GAT GGA TTG |
| NeoR-R |
| CTC GTC AAG AAG GCG ATA GAA |
| pCLN32PCRF2 |
| CTG ACT CTT AAT AAT GAA GGC TGC |
| pCLN32PCRR12 |
| TAT GAT GGA ACA CGT AAT GCG AGA |

Supplemental Table 2: Primers used for sequencing, targeting vector, rAAV production, and cell screening. Primers Listed from 5’ to 3’.

| | WT | | | | CLN3$^{\Delta ex7/8}$ | | | |
|---|---|---|---|---|---|---|---|---|
| | 24 | 30 | 36 | 42 | 24 | 30 | 36 | 42 |
| Fitbark | 2 | 9 | 10 | 2 | 5 | 9 | 11 | 4 |
| T-maze | 12 | 11 | 10 | 3 | 10 | 11 | 8 | 2 |

| | WT | | | | | CLN3$^{\Delta ex7/8}$ | | | | |
|---|---|---|---|---|---|---|---|---|---|---|
| | 24 | 30 | 36 | 42 | 48 | 24 | 30 | 36 | 42 | 48 |
| Gait | 11 | 12 | 11 | 1 | 2 | 10 | 10 | 10 | 2 | 2 |
| ERG | 4 | 13 | 12 | 2 | 2 | 8 | 12 | 11 | 3 | 3 |

Supplemental Table 3: Animal numbers by month for Fitbark , T-maze, Gait and ERG testing.
